# Mechanical coordination of counter-gradient growth maintains organ curvature in the apical hook

**DOI:** 10.64898/2026.01.31.703025

**Authors:** Sara Raggi, Hemamshu Ratnakaram, Adrien Heymans, Loitongbam Lorinda Devi, Özer Erguvan, Siamsa M. Doyle, François Jobert, Asal Atakhani, Sijia Liu, Manuel Petit, Jürgen Kleine-Vehn, Krzysztof Wabnik, Stéphane Verger, Stéphanie Robert

## Abstract

How growing tissues convert mechanical tension into signals that stabilize form remains a central question in morphogenesis. Curved organ shapes in plants arise from differential growth, yet how such curvature is actively maintained while organs continuously grow remains poorly understood. In etiolated seedlings, the apical hook provides a tractable model to dissect this process, as its curvature is stably maintained over extended periods despite ongoing cell expansion. Using quantitative imaging and computational modeling, we show that antagonistic growth gradients at apical and basal regions are both necessary and sufficient to maintain hook curvature, with cuticle integrity being critical for establishing these counter-gradients. Mechanical cues linked to cuticle structure, coupled with apoplastic reactive oxygen species (ROS), coordinate cellular growth anisotropy, and disruptions in cuticle biosynthesis trigger defective hook development. These findings reveal that the apical hook curve maintenance is not a simple switch between growth promotion and repression, but a highly dynamic, tightly regulated process where mechanical and biochemical signals coordinate organ-scale morphogenesis, fundamentally reshaping how we understand developmental growth.

## Introduction

Organ morphogenesis emerges from the spatiotemporal regulation of differential growth; whereby local domains of tissue undergo distinct rates and orientations of cell expansion to produce organ-specific geometries. Plant cell expansion is a mechanically constrained process governed by the interplay between intracellular turgor pressure and the viscoelastic properties of the cell wall, a polysaccharide network located outside the plasma membrane. The anisotropic mechanical properties of the wall, largely dictated by the spatial organization of cellulose microfibrils, the cross-linking of hemicelluloses, and the biochemical status of pectins, impose directional constraints that define the axis of cell elongation and ultimately organ shape ^1,2^. In plants, cells can be conceptualized as pressurized cylinders in which turgor-generated tensile stress exerts forces on the surrounding wall matrix. Irreversible expansion occurs only when the wall undergoes controlled yielding through enzymatic remodeling, mediated by wall-loosening agents such as expansins, xyloglucan endotransglycosylases, and other polysaccharide-modifying enzymes, concomitant with the incorporation of newly synthesized wall polymers to preserve structural integrity ^3–5^. Growth heterogeneity, such as when the cell elongates in one direction, is mediated by anisotropic cell wall loosening, cytoskeletal dynamics, and differential deposition of cell wall polymers, processes that are themselves tightly modulated by gradients of hormones such as auxin and cytokinin ^6,7^. At the tissue scale, the coordinated integration of these local growth patterns generates emergent mechanical stresses that create feedback to influence growth anisotropy and axis formation.

In the natural environment, seedlings germinate in darkness surrounded by soil, under physical conditions that can hinder proper emergence by inducing mechanical stress and physical damage. To cope with these challenges, dicotyledonous plants have evolved a transient inverted-U-shaped structure at the tip of the hypocotyl called the apical hook, which plays a vital protective role, shielding the shoot apical meristem and cotyledons during soil emergence. Apical hook development unfolds in three distinct phases: formation, maintenance, and opening ^8^. The transitions between these phases depend on precise control and modulation of growth rates, rendering the apical hook an exemplary system for investigating dynamic morphogenesis. Following germination, the hook is established through differential growth across the hypocotyl, with cells on one side elongating faster than those on the opposite side, driving hypocotyl arching ^8,9^. During the maintenance phase, differential growth continues at a stable rate, sustaining hook curvature. Hook straightening later occurs during the opening phase, when cells on the inner side elongate more rapidly than those on the outer side. The relationship between differential growth and morphogenesis is particularly evident during apical hook development, since gradients of cell expansion translate directly into visually perceptible bending of an organ. This makes the apical hook a valuable model for dissecting the complex interactions among signaling networks, hormone gradients, biomechanical cues, cell growth parameters, and organ morphogenesis ^10–15^. Mechanical feedback, mediated through wall stress and strain, integrates with gradients of hormones such as auxin, gibberellin and ethylene to coordinate asymmetric growth, ensuring robust formation and maintenance of the hook structure ^11,14,16^.

The outer cell walls of epidermal cells at the surface of plant above-ground organs including the apical hook, and transiently, the root meristem, are covered by a waxy, lipid-rich extracellular layer called the cuticle ^17,18^. This ancient structure emerged during the transition of plants to land, primarily to prevent dehydration and provide mechanical rigidity to early land plant structures ^19^. During evolution, the cuticle has since acquired additional roles beyond water retention, including protection against biotic and abiotic stresses, as well as acting as a core determinant of plant development ^17,20^.

Reactive oxygen species (ROS) are continuously produced in plants as by-products of metabolism but are also generated in a regulated manner by enzymatic systems such as NADPH oxidases in response to developmental and environmental cues ^21–23^. In addition to their signaling functions ^24^, ROS participate in cell wall remodeling and cuticle formation by promoting oxidative cross-linking of wall polymers and influencing cutin and wax assembly. Notably, cuticle-defective mutants often display altered ROS accumulation and signaling, highlighting a tight functional interplay between redox homeostasis, cell wall architecture, and cuticle integrity ^25,26^.

Here, we show using quantitative imaging and computational modeling that opposing growth gradients at the apical and basal halves of the apical hook are required to maintain curvature, with cuticle integrity being essential for establishing these gradients. Furthermore, we show that mechanical cues from the cuticle, together with apoplastic ROS, coordinate growth anisotropy, and disruptions in either system lead to defective hook development. These results highlight how mechanical and biochemical signals integrate to control organ-scale morphogenesis.

## Results

### Coordinated antagonistic growth dynamics maintain apical hook curvature

While the cellular mechanisms underlying hook formation and opening are well characterized ^12^, the processes sustaining curvature during the maintenance phase remain poorly understood. Previous studies have suggested that antagonistic growth gradients on opposite sides of the apical–basal midline preserve hook shape ^8,12,27–29^, but direct, quantitative evidence has been lacking. To address this, we generated a high-resolution spatial map of cell elongation across the apical hook. Time-lapse confocal imaging of *Arabidopsis thaliana* seedlings expressing the plasma membrane marker *p35S::PIP2A-GFP* was performed from ∼24 hours (T0) to ∼32 hours (T8) post-germination (Figure 1A). Changes in cell surface area were quantified using MorphoGraphX ^30^ (Figure 1B), and data from six seedlings across three biological replicates were integrated into a correspondence matrix capturing mean elongation (strain) rates at discrete positions along the hook (Figure 1C). Along the longitudinal axis of the hook, we refer to the cotyledon-proximal domain as apical and the hypocotyl-proximal domain as basal. Along the radial axis of the hook, curvature establishes opposing inner (concave) and outer (convex) domains. Analysis within this quantitative framework, with the orthogonal apical–basal and inner–outer axes, revealed the spatial pattern of growth distribution that underlies hook maintenance (Figure 1B and C). This approach provides the first high-resolution, quantitative map linking local cellular growth patterns to global organ curvature. In wild-type seedlings, strain rates differed between the inner and outer sides of the hook in both apical and basal regions. Maximal growth occurred in the inner–basal region, while substantial elongation was maintained on both the inner and outer sides of the apical region (Figure 1C and D). Quantitative analysis revealed two opposing anisotropic growth gradients across the hook. In the apical half, cell elongation was higher on the outer side, whereas in the basal half, elongation was higher on the inner side. Together, these apical and basal counter-gradients generate an antagonistic growth pattern that maintains hook curvature (Figure 1B and C).

**Figure 1.**
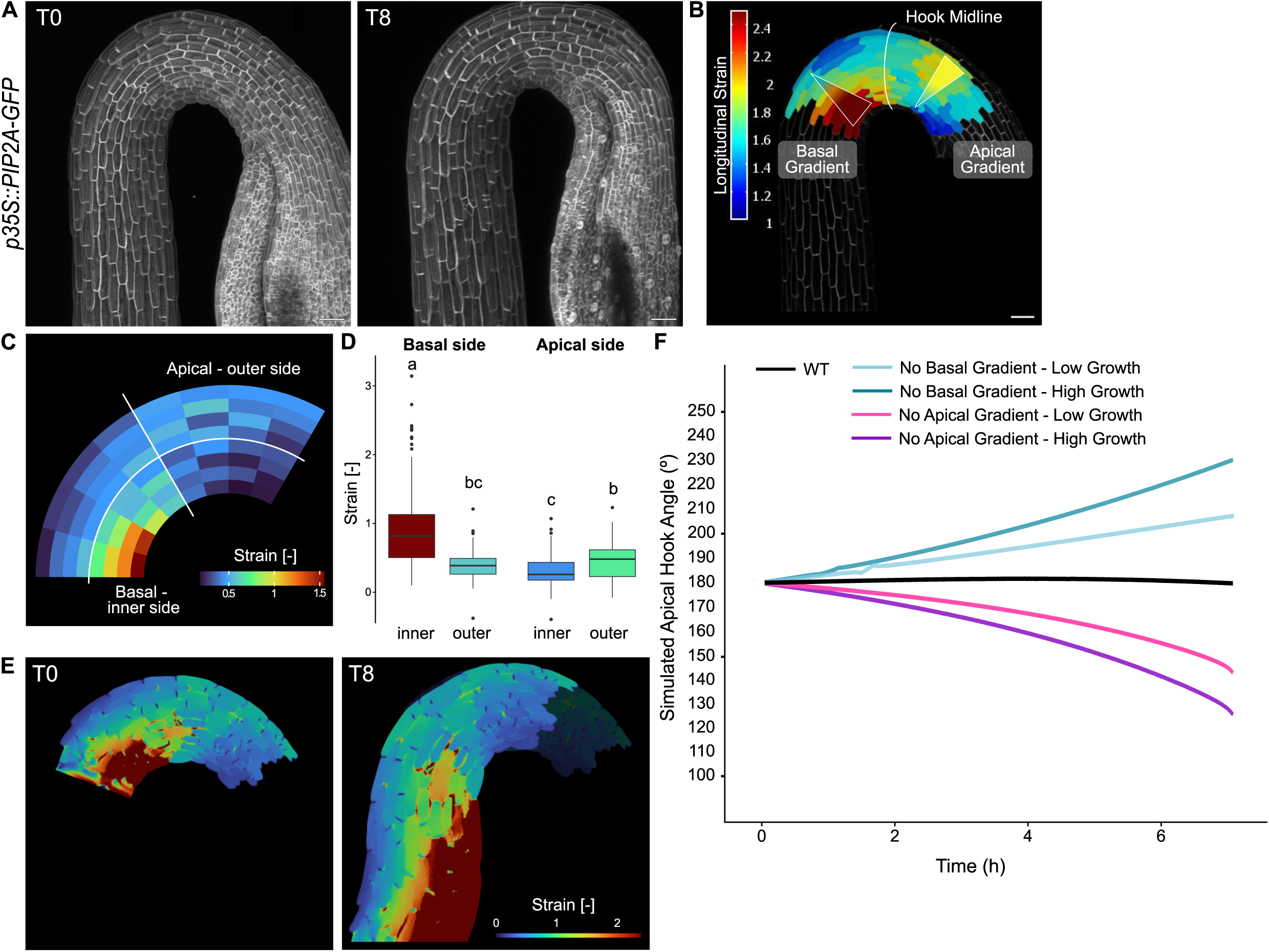
Characterization of differential growth patterns underlying apical hook maintenance in *Arabidopsis*. (A) Representative live confocal micrographs of dark-grown apical hooks of *Arabidopsis thaliana* expressing the plasma membrane marker *p35S::PIP2A-GFP* at approximately 24 h after germination (T0) and 8 h later (T8), illustrating hook maintenance during this developmental window. Scale bars represent 50 µm. (B) Schematic representation of the apical hook cell elongation domains and corresponding heatmap indicating the observed strain along the principal direction of growth observed from T0 to T8 in the representative *p35S::PIP2A-GFP* seedling pictured in (A). The apical–basal and inner–outer axis definitions represented in the schematic were used for growth quantification. Scale bar represents 50 µm. (C) Correspondence matrix of wild-type (*p35S::PIP2A-GFP*) longitudinal strain magnitudes, revealing two opposing gradients of differential growth, one in the apical region and one in the basal region, with higher expansion on the basal–inner and apical–outer sides. Data from six seedlings across three biological replicates were used to generate the matrix. (D) Boxplots of longitudinal strain for each domain shown in (C). ≥15 cells per domain were measured for each seedling. Different letters indicate statistically significant differences based on a Tukey post hoc test (α = 0.05). (E) Finite element model of a wild-type (*p35S::PIP2A-GFP*) apical hook mesh at T0 and the corresponding simulated geometry at T8, generated using average growth properties. (F) Simulated apical hook angle kinematics for a wild-type (*p35S::PIP2A-GFP*) mesh subjected to different growth scenarios: average growth (WT), average growth without the apical gradient and replaced by uniform low growth corresponding to the mean inner-domain elongation, average growth without the apical gradient and uniform high growth corresponding to the mean outer-domain elongation, average growth without the basal gradient and replaced by uniform high growth corresponding to the mean inner-domain elongation, or average growth without the basal gradient and replaced by uniform low growth corresponding to the mean outer-domain elongation.

Echoing previous reports ^8^, we observed an apparent apical–basal displacement of cells across the hook midline over 8 hours of growth, giving rise to a net directional “flow” (Figure S1A). Even though apical hook curvature appears to remain static during apical hook maintenance, the constant flow of cells requires a positional but not cell-dependent elongation program, where a single cell will show different rates of cell elongation over time, based on its relative position within the hook structure. These findings indicate that curvature maintenance arises not from static structural constraints but from the dynamic coordination of opposing growth fields across the hook.

To further investigate how spatial growth patterns translate into tissue-scale shape changes, we implemented a two-dimensional finite element model (FEM) of the apical hook, parameterized with cell-scale growth rates derived from our high-resolution dataset. Each cellular domain on the FEM mesh was assigned a distinct strain rate tensor, defining a unique growth scheme for every cell within the mesh, while strain-dependent growth shaped the extracellular domain. First, we simulated 8 hours of the hook maintenance phase using a reference mesh from a representative seedling (Figure 1E). Model benchmarking was performed by applying pooled growth parameters from seedlings expressing *p35S::PIP2A-GFP*, reproducing observed curvature dynamics (Figure 1E, T8). It should be noted that the 2D representations are not digital twins of the individual seedlings but rather provide a simplified framework for testing apical hook kinematics. Next, we simulated a perturbed local growth anisotropy by dissipating the elongation gradient in either the apical (Figure S1B) or basal (Figure S1C) domain. Dissipation of the apical gradient while preserving the basal gradient led to progressive hook opening, whereas dissipation of the basal gradient with retention of the apical gradient generated an over-hooking phenotype (Figure 1F). The magnitude of growth within the dissipated domain further modulated these effects: in the basal region, high uniform growth amplified over-hooking compared with low uniform growth (Figure 1F, S1C), while in the apical region, higher growth resulted a greater decrease in curvature (Figure 1F).

These simulations demonstrate that in a two-dimensional representation of the apical hook structure, a stable apical hook angle emerges from the dynamic equilibrium between two distinct and opposite counter-gradients of cell elongation that contribute antagonistically towards apical hook curvature. Together with our quantitative imaging, this computational modelling approach links cellular elongation to apical hook-scale morphogenesis.

### Cuticle integrity is essential for apical hook development

To better understand the factors coordinating the two opposing growth gradients on either side of the hook midline, we turned to genetic perturbations guided by our modeling predictions. Computational simulations indicated that disrupting the balance between apical and basal counter-gradients should primarily shorten the duration of curvature maintenance. We therefore focused on *hookback* (*hkb)* mutants previously isolated from a forward genetic screen for defects in auxin-dependent apical hook development ^31^. Among these, we identified two mutants, *hkb2* and *hkb4*, that displayed defects in apicl hook maintenance under control conditions, as revealed by kinematic analysis of apical hook angle over time ^9^. Both mutants displayed a similar phenotype, presenting a normal formation phase, shorter maintenance phase, and earlier and faster hook opening than the wild-type (Figure 2A). Whole genome sequencing revealed the presence of mutations in genes with roles in cuticle biosynthesis—the *CYTOCHROME P450, FAMILY 77, SUBFAMILY A, POLYPEPTIDE 4* (*CYP77A4)* gene in *hkb2* (G to A non-synonymous substitution at chromosome 5, position 1337420) and the *DEFECTIVE IN CUTICULAR RIDGES* (*DCR*) gene in *hkb4* (C-to-T non-synonymous substitution at chromosome 5, position 8077553)—suggesting a potential link between the cuticle and apical hook development. To confirm that the identified mutations were responsible for the apical hook development phenotype, we analyzed the hook angle kinematics phenotype of the insertional mutants *cyp77a4-4* and *dcr-2.* Both mutants displayed very similar phenotypes to *hkb2* and *hkb4*, with a shorter maintenance phase, and earlier and faster hook opening than the wild-type (Figure 2A), suggesting that the integrity of the cuticle layer might be important for proper apical hook development. We further examined apical hook development in *dcr-2* complemented with *pDCR::DCR-mCherry*, confirming that the developmental defects of this mutant were attributable to loss of DCR function (Figure S2A). We next investigated CYP and DCR expression patterns in *CYP77A4* and *DCR* mutant lines complemented with the functional fluorescent fusion constructs *pCYP77A4::CYP77A4-GFP* (in the *cyp77a4-3* background; Kawade et al. ^32^) and *pDCR::DCR-mCherry* (Figure S2B). Remarkably, we observed a gradual increase in CYP77A4 and DCR protein abundance over time, with relatively minimal accumulation at the early maintenance phase (22-24 hours post germination), greater protein expression at the maintenance phase (34-38 hours post germination) and highest expression at the late maintenance phase (46-52 hours post-germination), confirming a potential developmentally regulated role of CYP77A4 and DCR in apical hook morphogenesis.

**Figure 2.**
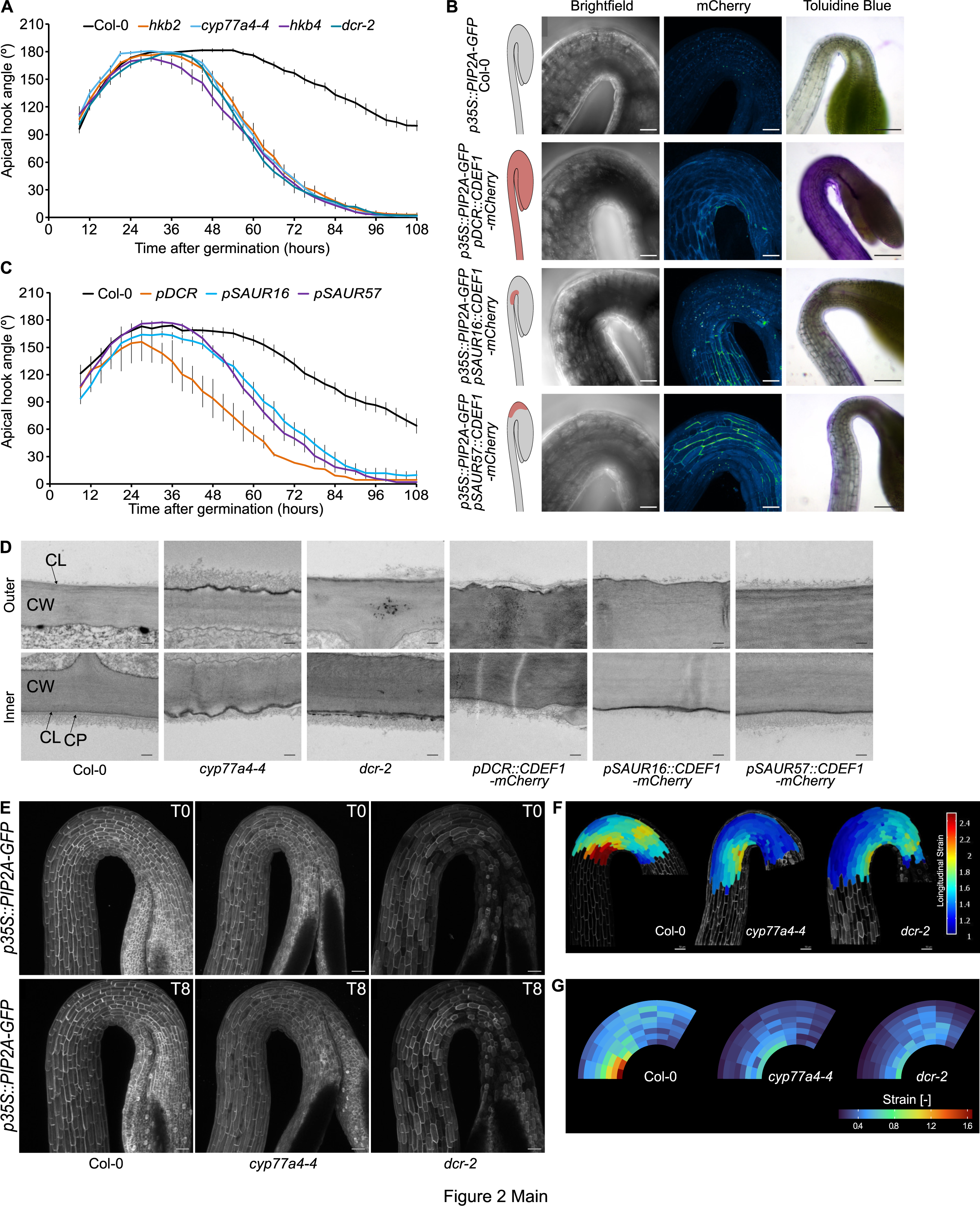
Cuticle integrity is essential for apical hook development. (A) Kinematics of apical hook angle in dark-grown Col-0 wild-type, *hkb2*, *cyp77a4-4*, *hkb4* and *dcr-2* as measured every 3 h, with 0 h as the time point of germination. The means of ≥19 seedlings per line are shown; error bars represent standard error of the mean. (B) Schematics of mCherry expression regions (red), representative live confocal micrographs on brightfield and mCherry channels and representative light micrographs after toluidine blue staining, of dark-grown upper hypocotyls/apical hooks of *p35S::PIP2A-GFP* in Col-0, *pDCR::CDEF1-mCherry*, *pSAUR16:: CDEF1-mCherry* and *pSAUR57:: CDEF1-mCherry* backgrounds in the maintenance phase (48 hours post-germination). Scale bars represent 50 µm in the confocal micrographs and 100 µm in the light micrographs. (C) Kinematics of apical hook angle in dark-grown seedlings of the lines shown in (B) as measured every 3 h, with 0 h as the time point of germination. The means of 19–24 seedlings per line are shown; error bars represent standard error of the mean. (D) Representative transmission electron micrographs of longitudinal sections of the inner and outer apical hook sides of the lines shown in (B) in the maintenance phase (32 hours post-germination). CW: cell wall; CL: cuticle layer; CP: cuticle proper. Scale bars represent 0.25 µm. (E) Representative live confocal micrographs of dark-grown apical hooks expressing the plasma membrane marker *p35S::PIP2A-GFP* in Col-0 wild-type, *cyp77a4-4* and *dcr-2* backgrounds at approximately 24 h after germination (T0) and 8 h later (T8). Scale bars represent 50 µm. (F) Longitudinal strain heatmap overlaid on representative T0 (24 hours post germination) micrographs indicating observed longitudinal strain along the principal direction of growth from T0 to T8 in apical hooks expressing *p35S::PIP2A-GFP* in Col-0 wild-type, *cyp77a4-4* and *dcr-2* backgrounds. One representative seedling per line is shown. Scale bar represents 50 µm. (G) Correspondence matrix of cell elongation strain rates in apical hooks expressing *p35S::PIP2A-GFP* in Col-0 wild-type, *cyp77a4-4* and *dcr-2* backgrounds. Data from four to six seedlings per line across three biological replicates were used to generate the matrix.

To establish that the defects in apical hook development are linked to cuticle integrity defects in the mutants, we analyzed the hook angle kinematics phenotype of four additional mutants for genes involved in cuticle biosynthesis or transport: *long-chain acyl-coA synthase 1* (*lacs1-1*), *glycerol-3-phosphate sn-2-acyltransferase 8* (*gpat8*), *atp-binding cassette g 11* (*abcg11-7*) and *bodyguard 1* (*bdg1-8*). While *lacs1-1* and *gpat8* exhibited a wild-type-like phenotype, *abcg11-7* and *bdg1-8* displayed a shorter maintenance phase and faster and earlier apical hook opening (Figure S2C), similar to the mutants affected in *CYP77A4* and *DCR*. We also assessed the cuticle integrity of these mutants in the apical region of the hypocotyl by performing toluidine blue staining on two-day-old seedlings ^33^. Toluidine blue is a cationic dye that associates with negatively charged cell wall constituents; under normal conditions, the cuticle prevents dye penetration, so that visible staining is diagnostic of increased cuticular permeability. Those mutants displaying altered apical hook development also showed strong toluidine blue staining in the apical hook region, while no staining was observed for Col-0, *lacs1-1* or *gpat8* (Figure S2D), reinforcing the hypothesis that cuticle integrity is necessary for proper apical hook development.

To further confirm that cuticle integrity plays a role during hook development, and to assess whether the cuticle present on either side of the apical hook might play differential roles, we locally expressed the *CUTICLE DESTRUCTING FACTOR 1* (*CDEF1*) gene of *Arabidopsis*, encoding an esterase that degrades cutin polyester ^34^. We fused *CDEF1* with the gene for the fluorescent label mCherry and expressed it under the control of the *DCR* promoter, which is active throughout the apical hook epidermis ^35,36^, and the *SMALL AUXIN UP-REGULATED RNA (SAUR) 16* and *57* promoters, as they are specifically expressed in the inner or outer side of the hook, respectively ^37^ (Figure 2B). First, we confirmed using confocal microscopy that CDEF1-mCherry was expressed across the apical hook when driven by *pDCR* and enriched in the inner or outer sides when driven by the *pSAUR16* and *57* promoters respectively (Figure 2B). We then verified the extent of cuticle integrity loss induced by CDEF1 activity with toluidine blue staining and found that the pattern of staining was comparable to the domain of CDEF1 expression (Figure 2B). Apical hook development in the *CDEF1*-expressing lines was then analyzed over time (Figure 2C). Similarly to the cuticle biosynthesis mutants *cyp77a4-4* and *dcr-2*, the *CDEF1-*expressing lines displayed a shorter maintenance phase and earlier and faster apical hook opening compared to the wild-type control (Figure 2C). In particular, the degradation of the cuticle across the whole apical hook region (under *pDCR*-driven expression) induces even earlier opening than observed in *cyp77a4-4*, *dcr-2* or the *pSAUR*-driven expressors of *CDEF1*, most probably due to the high activity of the esterase. No major differences could be observed between the lines with loss of cuticle integrity in the inner or outer hook side. Overall, our results clearly indicate that cuticle integrity on both sides of the hook is essential for proper apical hook morphogenesis, and that local loss of cuticle integrity likely acts in a non-cell autonomous fashion.

To analyze cuticle ultrastructure in the apical hook of our cuticle-affected lines, we performed transmission electron microscopy (TEM) in longitudinal sections of hooks in the maintenance phase (approximately 30 to 36 hours post-germination) (Figure 2D). In the control line, the cuticular layer of the outer side of the hook is adjacent to the cell wall and appears as an opaque diffused longitudinal deposition. On the inner side, the cuticular layer appears thin and compact, coated by a thicker, electron-translucent layer of cuticle proper. Thus, the cuticle structure differs between the outer and inner side of the hook during the maintenance phase, suggesting that its status changes dynamically, likely in coordination with the cell growth rate. In the *cyp77a4-4* and *dcr-2* mutants and most of the *CDEF1*-expressing lines, we revealed an abundant electron opaque deposition on both the outer and inner sides. These results show qualitative disruptions in the deposition and/or organisation of the cuticular layer in these lines. Taken together, the results confirm that the cuticle biosynthesis mutants carry a defect in cuticle ultrastructure and permeability and establish this defect as the cause of the observed defect in apical hook morphogenesis.

### Cuticle integrity regulates coordinated antagonistic growth dynamics

To further understand the cuticle’s role in apical hook morphogenesis, we analyzed cell size in the apical hook region in Col-0 and the cuticle biosynthesis mutants stained with propidium iodide (PI) cell wall dye at 24 hours after germination. We observed statistically significant reductions of 11% and 24% in mean cell length in the apical hook region of *cyp77a4-4* and *dcr-2* mutant backgrounds, respectively, compared to the wild-type Col-0 background (Figure S2E), implicating cuticle involvement in the regulation of cell and organ geometry. These data are suggesting a possible reduction in cell elongation rates or increased cell division frequency in *cyp77a4-4* and *dcr-2* compared to Col-0. To decipher between these two possibilities, we used the speed of light-induced apical hook opening as a readout of cell elongation rates on the inner side of the apical hook ^38^. We observed that the mutants *cyp77a4-4* and *dcr-2* displayed a statistically significant reduction in the rate of light-induced apical hook opening compared to Col-0 (Figure S2F), supporting the idea of impaired cell elongation in the mutants.

We then performed time-lapse confocal imaging on the *p35S::PIP2A-GFP* expressing lines in the wild-type Col-0, *cyp77a4-4* and *dcr-2* backgrounds at 24 hours (T0) and 32 hours (T8) after germination to directly observe the magnitude and distribution of cell expansion (Figure 2E and F). Data from 4-6 seedlings per line across three biological replicates were integrated into a correspondence matrix to spatially map cell expansion during apical hook maintenance in the apical hooks of *cyp77a4-4* and *dcr-2* (Figure 2G). In these cuticle defective mutants, we observed a disruption in the counter gradient system compared to the Col-0 wild-type (Figure 2F and G). Importantly, the apical counter-gradient was nearly abolished in *cyp77a4-4* and *dcr-2* during apical hook maintenance, while the basal gradient appeared to be less steep in comparison to that of the wild-type Col-0 (Figure S2G), suggesting that cuticle integrity might be involved in regulating the counter-gradients required for apical hook maintenance.

Using the FEM framework we previously established (Figure 1E, F), we simulated the development of representative *cyp77a4-4* and *dcr-2* apical hooks constructed using data from the time-lapse experiments to translate the quantitative measurements of cell elongation to insights into the morphodynamics of the mutant apical hooks (Figure 3A, B). To understand if the mutants disrupted counter-gradients are sufficient to explain their hook development phenotype, we simulated the development of a wild-type reference mesh with each mutant’s cell elongation patterns (Figure 3C). Under both the *cyp77a4-4* and *dcr-2* growth regimes, the wild-type reference meshes exhibited reduced hook angles at the end of the simulation compared with the control wild-type growth regime, which maintained a stable curvature throughout the simulation (Figure 3B, C); this suggests that the disrupted counter-gradients in *cyp77a4-4* and *dcr-2* sufficiently account for their defects in apical hook development. When we computationally applied the growth rates of each mutant onto the mutants’ own cell geometries (Figure 3D), we observed a reduction in hook angle compared to the control wild-type growth regime, further supporting the idea that growth patterns sufficiently account for the mutant apical hook development phenotype. Applying wild-type growth patterns to the *dcr-2* geometry restored apical hook maintenance (Figure 3D); the *cyp77a4-4* geometry, however, displayed an exaggeration of hook curvature under a wild-type growth regime (Figure 3D), suggesting that initial tissue geometry also contributes to apical hook development dynamics. These simulations indicate that the absence of counter-gradient growth provides a strong morphomechanical basis for the premature hook-opening phenotype observed in cuticle-defective mutants.

**Figure 3.**
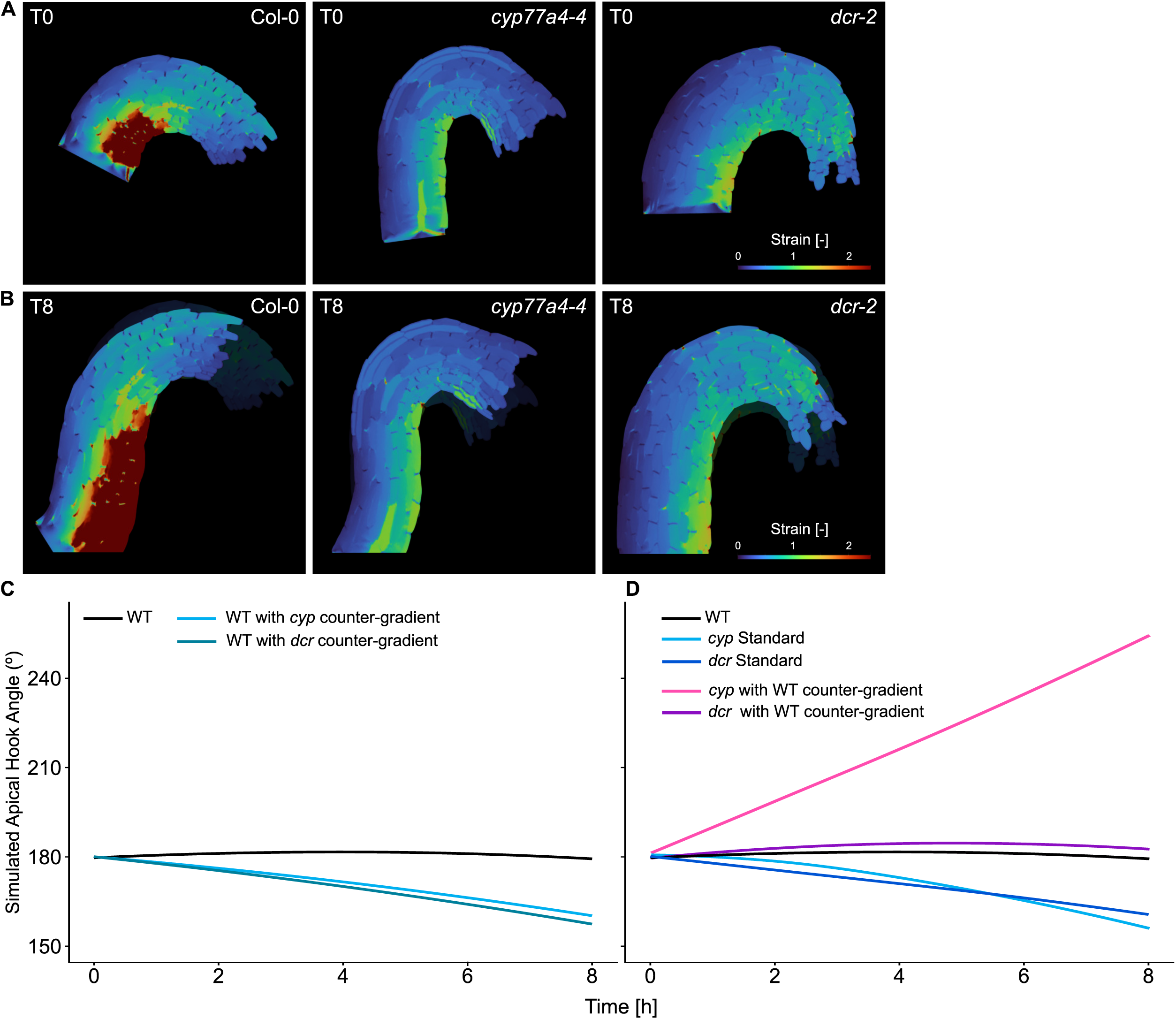
Altered growth properties and counter-gradients affect apical hook kinematics. (A-B) Finite element models of apical hooks of wild-type Col-0, *cyp77a4-4* and *dcr-2*, showing the initial mesh at T0 (A) and the corresponding simulated geometry after 8 h of development (T8) (B). Simulations were generated using the average growth properties of the respective genotypes. (C) Kinematics of WT apical hook simulated using either WT, *cyp77a4-4*, or *dcr-2* counter-gradient profiles. (D) Kinematics of WT, *cyp77a4-4*, and *dcr-2* apical hooks simulated using the growth properties of their respective genotypes, as well as *cyp77a4-4* and *dcr-2* apical hook simulated with WT counter-gradient profiles.

### Mechanical cues link cuticle integrity to the control of counter-gradients during apical hook maintenance

Cell expansion is driven by the biomechanical feedback between turgor pressure and a structurally anisotropic cell wall, the extension and extensibility of which is directed and dynamically regulated by mechanical stress patterns and regulators of cell wall structure and composition ^6,39–43^. The observed changes in differential growth patterns at the apical hook in the cuticle biosynthesis mutants *cyp77a4-4* and *dcr-2* prompted us to assess the potential involvement of the cuticle in regulating the biomechanics of cell growth. We first visualized subcellular fluorescent markers known to be responsive to mechanical stress patterns. Sites of tissue compression in the shoot apical meristem boundaries display deformed nuclei consistent with the extent of organ growth ^44^, demonstrating that the shape of the nuclear envelope reflects tissue-scale mechanical stress patterns. Nuclear envelope shape can be quantified as the ratio between height and width (aspect ratio) of an idealized ellipse that best fits the observed nuclear boundary; an aspect ratio value of 1 indicates a perfect circle, with higher values indicating an increasingly deformed nuclear shape. We observed the nuclear envelope marker SAD1/UNC-84 DOMAIN PROTEIN 1 (*p35S*::*SUN1-YFP*) in apical hooks of the Col-0 wild-type, *cyp77a4-4* and *dcr-*2 backgrounds and quantified nuclear shape. The mean aspect ratio of nuclei in the apical hooks of all lines was significantly different between the outer and inner hook sides (Figure 4A-D). The Col-0 mean nuclear aspect ratio was 15% higher in the outer compared to the inner side, while in *cyp77a4-4*, the mean aspect ratio of nuclei was only 12% higher on the outside compared to the inside (Figure 4A, B). Furthermore, the mean aspect ratio of nuclei in *cyp77a4-4* was lower by 8% and 11% on the inside and outside respectively compared to the corresponding Col-0 nuclei. A very similar trend was also observed in the experiments with *dcr-2* (Figure 4C, D). Overall, we observed that increased cuticle permeability is associated with rounder nuclei on average, reflecting altered mechanical stress patterns in the mutants.

**Figure 4.**
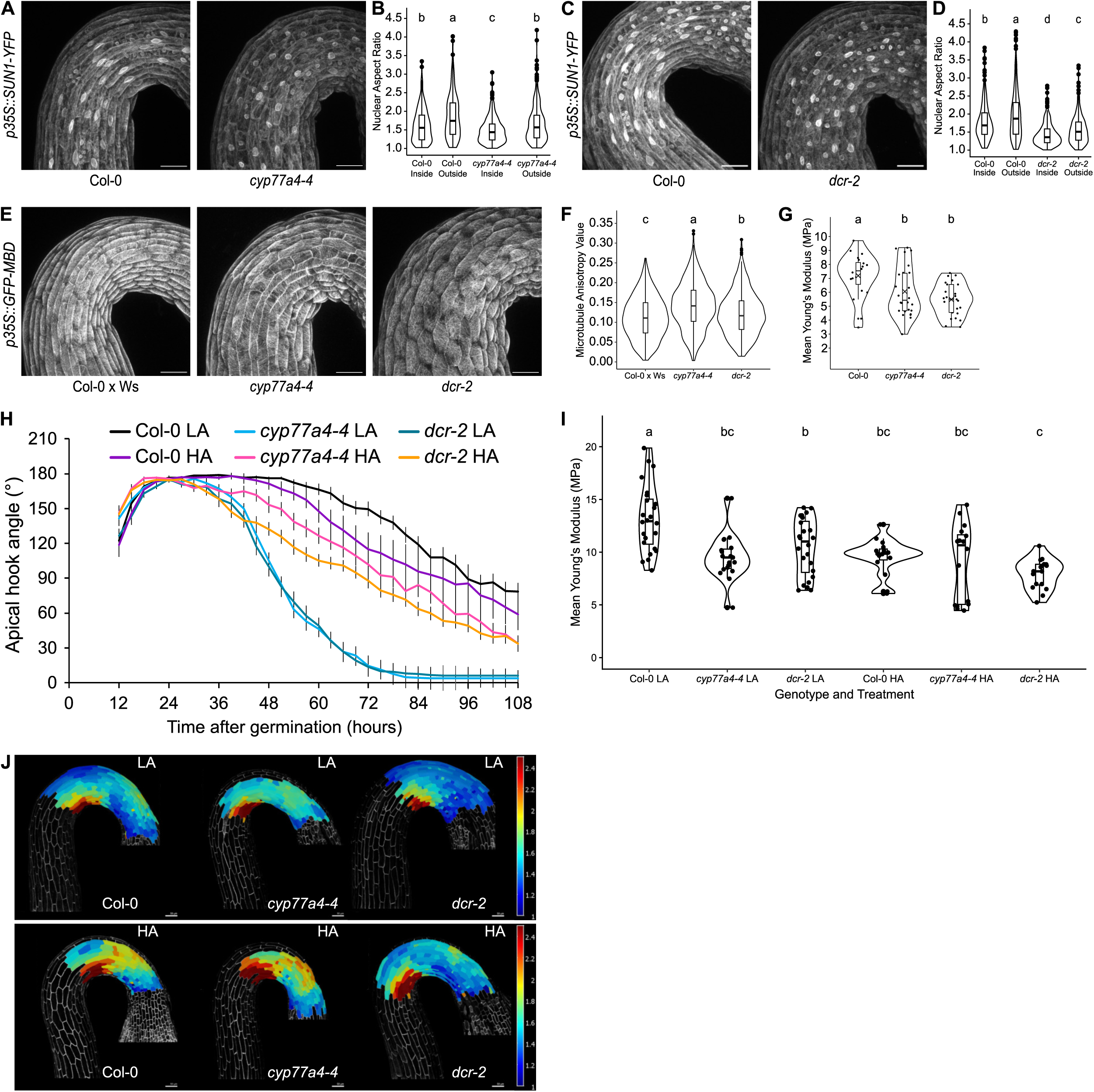
Mechanical cues link cuticle integrity to apical hook development. (A-D) Representative live confocal micrographs of dark-grown apical hooks of *p35S::SUN1-YFP* in Col-0 wild-type and *cyp77a4-4* backgrounds (A) or *dcr-2* backgrounds (C) and violin plots of nuclear aspect ratios (major axis length/minor axis length) of epidermal cell nuclei in the inside or outside half of apical hooks of *p35S::SUN1-YFP* in Col-0 wild-type and *cyp77a4-4* backgrounds (B) or *dcr-2* backgrounds (D) at approximately 48 h after germination, see representative images in (A) and (C). Scale bars represent 50 µm. ≥45 nuclei were measured for each of ≥9 seedlings per line across ≥3 biological replicates. Different letters indicate statistically significant differences based on one-way ANOVA and Tukey’s *post-hoc* tests (α = 0.05). (E) Representative live confocal micrographs of dark-grown apical hooks of *p35S::GFP-MBD* in Col-0 x Ws-4 wild-type, *cyp77a4-4* and *dcr-2* backgrounds. Scale bars represent 50 µm. (F) Violin plots of microtubule anisotropy values of epidermal cells in apical hooks expressing the nuclear envelope marker *p35S::GFP-MBD* in Col-0 x Ws-4 wild-type, *cyp77a4-4* and *dcr-2* backgrounds at approximately 24 h after germination, see representative images in (E). ≥40 cells were measured for each of ≥10 seedlings per line across ≥3 biological replicates. Different letters indicate statistically significant differences based on one-way ANOVA followed by Tukey’s *post-hoc* tests (α = 0.05). (G) Violin plots of Young’s Modulus (cell wall indentation stiffness) of epidermal cells in basal regions of dark-grown hypocotyls of *p35S::PIP2A-GFP* in Col-0, *cyp77a4-4* and *dcr-2* backgrounds, see representative images in Fig. S4A. 3 cells were measured for each of 3 seedlings per line across ≥3 biological replicates. Different letters indicate statistically significant differences based on one-way ANOVA and Tukey’s *post-hoc* tests (α = 0.05). (H) Kinematics of apical hook angle in dark-grown Col-0 wild-type, *cyp77a4-4* and *dcr-2* germinated and grown on 0.8% w/v agar medium (low-agar, LA) or 2.5% w/v agar medium (high-agar, HA) as measured every 3 h, with 0 h as the time point of germination. The means of ≥18 seedlings per line are shown; error bars represent standard error of the mean. (I) Violin plots of Young’s Modulus (cell wall indentation stiffness) of epidermal cells in basal regions of dark-grown hypocotyls of *p35S::PIP2A-GFP* in Col-0, *cyp77a4-4* and *dcr-2* backgrounds grown on LA and HA. 5–6 cells were measured for each of 5 seedlings per line, per growth condition. Different letters indicate statistically significant differences based on one-way ANOVA and Kruskal-Wallis tests, followed by Dunn’s test for pairwise comparisons (α = 0.05). (J) Heatmaps of longitudinal strain along the principal axis of growth observed in a representative seedling from T0 to T8 in apical hooks expressing *p35S::PIP2A-GFP* in Col-0 wild-type, *cyp77a4-4* and *dcr-2* backgrounds grown on LA and HA. One representative seedling per line and growth condition out of two biological replicates is shown. The heatmaps are each superimposed on the T0 micrograph of one representative seedling. Scale bar represents 50 µm.

Cortical microtubule organization is known to respond to tissue-scale mechanical stress patterns, aligning with the direction of maximal tensile stress ^45–48^. We imaged the outer face of epidermal cells in apical hooks of the cortical microtubule marker line *p35S::GFP-MBD* in Col-0 x Ws-4, *cyp77a4-4* and *dcr2* backgrounds and analyzed microtubule organization using the SurfCut2, Segmentation4FTBatch, FibrilTool, Cell_Aspect_Ratio and Angle2Cell ImageJ macros ^49,50^. Compared to the wild-type background, the mutants *cyp77a4-4* and *dcr-2* displayed statistically significant hyper-alignment of cortical microtubules (Figure 4E–F), with approximately 26% and 8% higher mean anisotropy than the wild-type background, respectively, strengthening the idea that cuticle integrity plays a role in regulating and redistributing mechanical stress patterns.

To assess the biomechanical impact of cuticle permeability, we used atomic force microscopy (AFM) to directly quantify cell wall indentation stiffness in Col-0 and the cuticle mutants. Since our cell elongation data was recorded in the *p35S::PIP2A-GFP* background, we used the same lines for the AFM experiments. Technical difficulties made it unfeasible to measure apparent cell wall stiffness directly in the apical hook. However, we observed that cells at the base of the hypocotyl also displayed significantly shorter and more anisotropic cells in *cyp77a4-4* and *dcr-2*, with a ∼20% reduction in cell length in the mutants, compared to the wild-type background (Figure S3A–B), making this region relevant for indentation stiffness measurements. We observed that the mean cell wall indentation stiffness (Young’s modulus) in the basal region of Col-0 background hypocotyls was significantly different to that of the mutants, being around 12% higher than in *cyp77a4-4* and *dcr-2* backgrounds (Figure 4G), demonstrating that a disruption of cuticle integrity results in changes in cell wall biomechanics, directly and/or through a putative signal associated with cuticle integrity.

Previous studies have demonstrated that reducing the water potential of solid growth media by increasing the concentration of agar results in a reduction of epidermal tension and apparent cell wall stiffness ^45^. In previous studies, increased agar concentration has rescued apical hook developmental defects in cell wall integrity-affected mutants ^51^. We tested the relationship between cuticle integrity and cell wall biomechanics when faced with changes in water availability by monitoring hook development in the wild-type Col-0, *cyp77a4-4* and *dcr-2* when grown on medium with high concentration of agar (2.5% w/v, hereafter referred to as high-agar or HA), compared to medium with our usual agar concentration (0.8% w/v, also referred to as low-agar or LA) (Figure 4H). Col-0 apical hooks opened slightly earlier on HA when compared to LA. Interestingly, the apical hook kinematics of the cuticle-defective mutants *cyp77a4-4 and dcr-2* were modified on HA compared to LA, opening more slowly and shifting towards wild-type-like apical hook development. Importantly, growth on HA did not rescue the cuticle integrity defects, as seen with toluidine blue staining (Figure S3C), suggesting that the observed change in apical hook kinematics in *cyp77a4-4* and *dcr-2* is independent of cuticle integrity. When grown on HA medium, hypocotyls of *p35S::PIP2A-GFP* in the Col-0 background displayed a significant 27% reduction in mean indentation stiffness of the epidermal cell walls measured using AFM, bringing the apparent stiffness to a level statistically indistinguishable from the *cyp77a4-4* and *dcr-2* backgrounds grown on HA medium (Figure 4I). The apparent indentation stiffness remained unchanged in the *cyp77a4-4* and *dcr-2* mutants grown on HA compared to LA (Figure 4I); however, the shift in apical hook kinematics observed in *cyp77a4-4* and *dcr-2* (Figure 4H) suggests that the mutants are sensitive to changes in water potential. Taken together, the results suggest that growth on LA is associated with distinct mechanical cues (that are absent when grown on HA) that in turn disrupt apical hook maintenance in the mutants *cyp77a4-4* and *dcr-2.* Analysis of the differential cell growth patterns over 8 hours (at 24 hours (T0) and 32 hours (T8) after germination) in apical hooks of *p35S::PIP2A-GFP* in Col-0, *cyp77a4-4* and *dcr-2* backgrounds grown on HA revealed an apparent restoration of the counter-gradients in the cuticle biosynthesis mutants compared to when grown on LA (Figure S3D-E and Figure 4J). Taken together, the above results suggest that mechanical cues associated with cuticle integrity orchestrate the counter-gradients required for apical hook development.

### Reactive oxygen species (ROS) link biomechanical cues with cuticle permeability in the control of counter-gradients during apical hook maintenance

Increased cuticle permeability has been shown to be associated with elevated production of reactive oxygen species (ROS) in the apoplast ^25,26,52^ and ROS accumulation has been observed in the apical hook ^53^. Using 3,3’-diaminobenzidine (DAB) staining ^54^, we observed elevated apoplastic hydrogen peroxide levels in the apical hook of the mutants *dcr-2* and *cyp77a4-4* when grown on LA, compared to the Col-0 wild-type (Figure 5A). In order to better understand this ROS accumulation and the mechanical stress patterns observed in *cyp77a4-4* and *dcr-2*, we also performed DAB staining in HA growth conditions. Interestingly, apoplastic ROS levels were reduced in *cyp77a4-4* and *dcr2* when grown on HA compared to LA (Figure 5A), indicating that mechanical cues link cuticle permeability status to apoplastic hydrogen peroxide levels in the apical hook during the maintenance phase.

**Figure 5.**
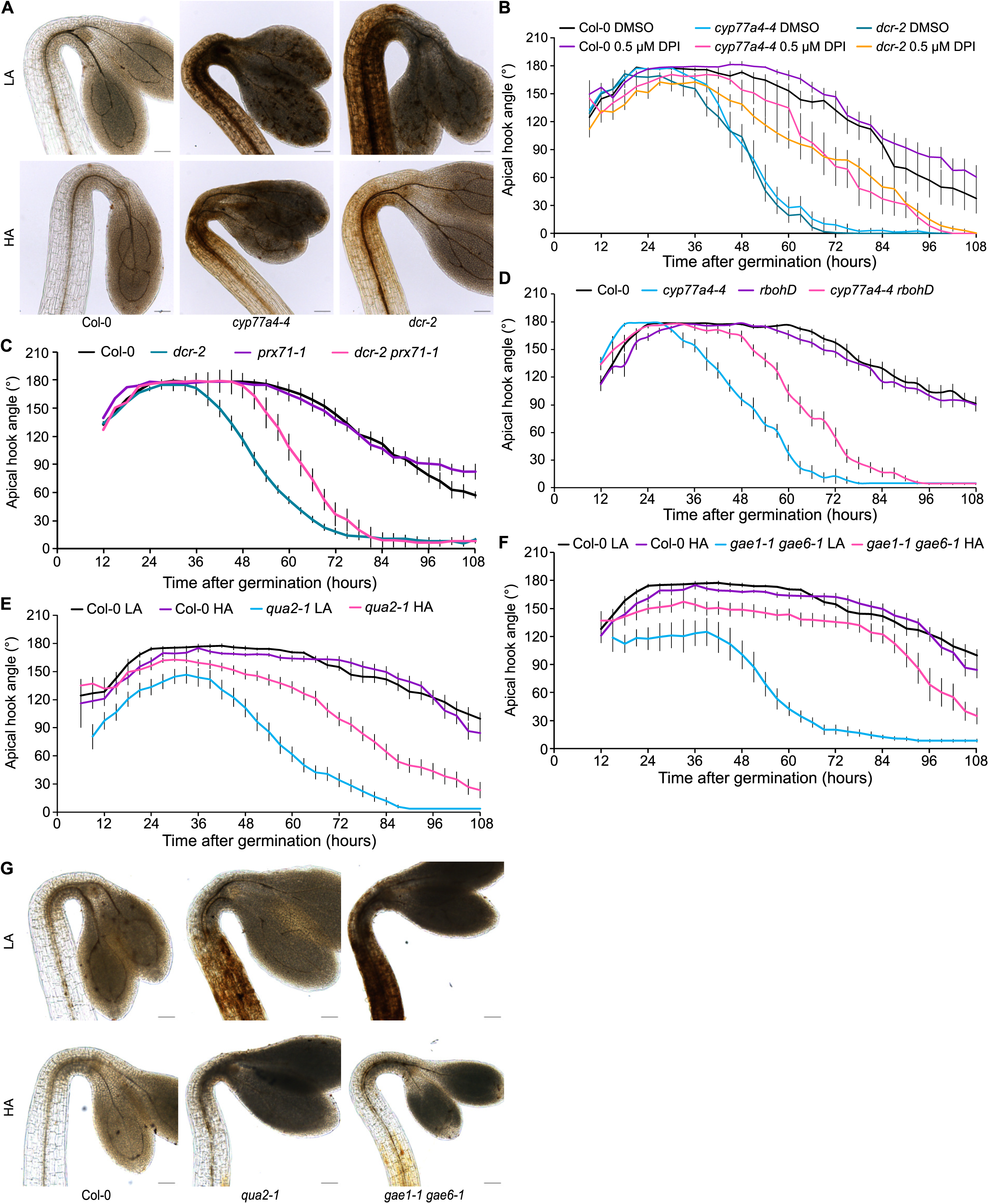
Reactive oxygen species (ROS) link biomechanical cues with cuticle permeability during apical hook maintenance. (A) Representative light micrographs of dark-grown upper hypocotyls and apical hooks of Col-0, *cyp77a4-4* and *dcr-2* grown on medium with low-agar (LA, 0.8% w/v agar) and high-agar (HA, 2.5% w/v agar) in the maintenance phase (48 hours post-germination) after DAB staining. 5 to 8 seedlings were analyzed for DAB staining over 3 biological replicates. Scale bars represent 100 µm. (B) Kinematics of apical hook angle in dark-grown Col-0 wild-type, *cyp77a4-4* and *dcr-2* germinated and grown on control medium (mock-supplemented with DMSO) or medium supplemented with 0.5 µM DPI as measured every 3 h, with 0 h as the time point of germination. The means of 8–14 seedlings per line are shown; error bars represent standard error of the mean. (C-D) Kinematics of apical hook angle in dark-grown Col-0 wild-type, *cyp77a4-4*, and *prx71-1* and *cyp77a4-4 prx71-1* (C) or *rbohD* and *cyp77a4-4 rbohD* (D) as measured every 3 h, with 0 h as the time point of germination. The means of 16–18 seedlings per line are shown; error bars represent standard error of the mean. (E-F) Kinematics of apical hook angle in dark-grown Col-0 wild-type and *qua2-1* (E) or g*ae1-1 gae6-1* (F) germinated and grown on 0.8% w/v agar medium (low-agar, LA) or 2.5% w/v agar medium (high-agar, HA) as measured every 3 h, with 0 h as the time point of germination. The means of 10–20 seedlings per line, per growth condition are shown; error bars represent standard error of the mean. (G) Representative light micrographs of dark-grown upper hypocotyls and apical hooks of Col-0, *qua2-1* and *gae1-1 gae6-1* grown in the dark on LA or HA in the maintenance phase (48 hours post-germination) after DAB staining. 5 to 8 seedlings were analyzed for DAB staining over 3 biological replicates. Scale bars represent 100 µm.

To dissect the relationship between elevated apoplastic ROS accumulation and coordinated growth necessary for apical hook maintenance, we utilized both pharmacological and genetic approaches. Diphenyleneiodonium (DPI) treatment inhibits the activity of NADPH oxidases and thereby ROS production ^55^. Upon treatment with DPI, we confirmed a reduction in apoplastic hydrogen peroxide levels in *cyp77a4-4* and *dcr-2* and via DAB staining (Figure S4B) and observed a prolongation of the maintenance phase in Col-0 and both mutants compared to mock treatment (Figure 5B). We also observed that the reduction in apoplastic ROS accumulation under DPI treatment did not restore the cuticle permeability defect in the mutants (Figure S4A). The prolongation of apical hook maintenance upon DPI treatment suggests a role for NADPH oxidase-mediated ROS production in regulating coordinated cell growth patterns. To confirm this, we analyzed apical hook development in plants overexpressing *PEROXIDASE71* (*p35S::PRX71*), which show increased ROS accumulation ^56^, as well as in the *peroxidase71-1* (*prx71-1*) and *respiratory burst oxidase homolog D (rbohD*) mutants, both of which are known to exhibit reduced ROS accumulation in response to isoxaben treatment ^56^. Notably, the *p35S::PRX71* line displayed a shortened maintenance phase and premature apical hook opening compared with the wild-type Col-0 (Figure S4C), further supporting a role for ROS in the regulation of apical hook development. The line *prx71-1* displayed similar hook development kinematics to Col-0 (Figure 5C) while the *rbohD* mutant had a slightly premature onset of hook opening compared the wild-type Col-0 (Figure 5D). To assess the effect of genetic inhibition of ROS production in the cuticle mutants, we crossed *prx71-1* and *rbohD* with *dcr-2* and *cyp77a4-4* respectively. The double mutant *dcr-2 prx71-1* displayed a prolongation of the maintenance phase compared to *dcr-2* (Figure 5C). We were unable to obtain double-homozygous *dcr-2 rbohD* lines, most likely due to embryo lethality issue in the double homozygote line. Interestingly, the double mutant *cyp77a4-4 rbohD* also showed a prolonged maintenance phase compared to the single mutant *cyp77a4-4* (Figure 5D). Taken together, these results suggest that the premature apical hook opening phenotype observed in *dcr-2* and *cyp77a4-4* is caused by elevated ROS accumulation.

### Biomechanical cues also link cell wall integrity with apoplastic ROS levels during apical hook development

Our results link apoplastic ROS levels to biomechanical cues associated with cuticle permeability. To determine whether apoplastic ROS also respond to general biomechanical signals, such as those arising from the cell wall, we measured apical hook kinematics and apoplastic ROS levels in LA and HA growth conditions using cell wall-affected mutant lines previously characterized for defective hook maintenance, which can be rescued under HA conditions ^51^. We thus monitored apical hook development over time in the mutants *quasimodo2-1* (*qua2-1*) and *glucuronate 4-epimerase1* and *6* (*gae1-1 gae6-1*), which are defective in pectin biosynthesis. In these mutants, defective apical hook maintenance results from reduced anisotropic growth on the outer side of the hook ^51^. Consistent with previous reports, both mutants exhibited defective hook maintenance under LA conditions, and the defect was partially rescued under HA conditions (Figures 5E–F). We next assessed apoplastic ROS accumulation in these mutants. During the maintenance phase (48 hours post-germination), both mutants grown on LA exhibited elevated apoplastic ROS levels compared to the wild-type Col-0 grown on LA (Figure 5G). In HA conditions, however, ROS accumulation was reduced in *qua2-1* and *gae1-1 gae6-1*, (Figure 5G) suggesting a complex feedback mechanism between mechanochemical properties of the cell wall, growth repartition and ROS accumulation.

Together, our work demonstrates that organ morphogenesis is not merely driven by growth promotion or repression, but is a highly dynamic, tightly regulated process where mechanical signals such as the ones issued from cuticle properties, and ROS integration coordinate development, shifting the paradigm for how we understand morphogenesis.

## Discussion

Our study identifies a counter-anisotropic growth gradient as the core mechanism sustaining apical hook curvature during the maintenance phase. While differential growth has long been implicated in hook morphogenesis ^57^, curvature maintenance has often been viewed as a passive or mechanically stable state. Our quantitative growth mapping instead reveals that maintenance is an active process, requiring the continuous coordination of opposing elongation fields. This counter-gradient represents a conceptual advance over classical models that invoked simple growth differentials to explain hook shape. Although antagonistic growth zones were previously proposed based on kinematic and physiological observations ^8,12,27–29^, direct quantitative evidence at cellular resolution was lacking. By integrating time-resolved imaging with mechanical modeling, we demonstrate that these antagonistic growth patterns are both necessary and sufficient to maintain curvature. Selective dissipation of growth anisotropy in either apical (such as in *cyp77a4-4* and *dcr-2* mutants) or basal domains was sufficient to destabilize hook shape, driving premature opening or exaggerated curvature, respectively. Our findings further suggest that transitions between hook formation, maintenance, and opening are regulated by rebalancing pre-existing growth fields rather than by abrupt switches in growth programs. This transition most likely involves complex hormonal crosstalk ^58^. Such a mechanism provides robustness while preserving developmental flexibility, allowing rapid modulation of hook curvature in response to environmental cues. More broadly, counter-gradient growth dynamics may represent a general principle of plant morphogenesis, enabling the stabilization of curved or asymmetric forms while maintaining the capacity for rapid shape change in diverse developmental contexts.

Although traditionally regarded as a protective barrier (such as in the root cap ^18^, the plant cuticle has increasingly been implicated in the regulation of growth and tissue mechanics ^20^. Our findings reveal a temporal uncoupling between the establishment and maintenance of differential growth during apical hook development. While the cuticle-defective *cyp77a4-4* and *dcr-2* mutants underwent a normal formation phase, indicating that cuticle integrity is dispensable for the initial establishment of growth asymmetry, disruption of cuticle integrity at later stages selectively impaired hook maintenance. This loss of curvature correlated with altered antagonistic growth gradients, demonstrating that sustained differential growth requires an intact cuticle. Consistently, the accumulation of CYP77A4 and DCR during the maintenance phase indicates that CYP77A4- and DCR-dependent cuticle biosynthesis is tightly and developmentally regulated. Together, these results uncover a phase-specific role for the cuticle, positioning it as a key mechanical and regulatory component that preserves growth asymmetry after it has been established, rather than contributing to its initial formation. In line with previous observations that cuticle-defective mutants display reduced cell elongation and elevated CYCB1 activity in etiolated hypocotyls ^36,59^, we observed a general reduction in cell expansion rates and anisotropy in cuticle-deficient apical hooks. These effects lead to a pronounced attenuation of the antagonistic growth gradients required for curvature maintenance. Mechanical simulations further support the conclusion that altered growth dynamics, rather than cellular and tissue geometry alone, are sufficient to destabilize hook curvature, highlighting the importance of cuticle integrity for coordinating spatial growth patterns. Beyond the apical hook, cuticle-mediated developmental regulation has been implicated in other contexts. For example, in petals and embryo development, cuticle integrity contributes to proper tissue patterning and prevents undesired organ fusion, consistent with a role in coordinating growth across tissues ^60,61^. Together with studies of expanding organs such as tomato fruit, which demonstrate that cuticle mechanical properties are developmentally regulated and spatially heterogeneous ^62–65^, these findings support a broader view of the cuticle as a structural component that contributes to the mechanical framework underlying coordinated growth and morphogenesis across plant development.

Perturbations in cell wall integrity trigger ROS production as part of a tightly regulated compensatory and signaling response. Altered wall architecture, such as reduced cellulose or pectin integrity, is sensed by cell wall integrity–monitoring pathways involving receptor-like kinases such as THESEUS1 and FERONIA, which initiate downstream ROS-generating mechanisms ^66,67^. ROS production occurs through both NADPH oxidases and apoplastic Class III peroxidases (CIII Prxs), providing spatially localized oxidative signals that reinforce wall structure and modulate growth ^68^. Notably, it has been demonstrated that the peroxidase PRX71 is upregulated in response to cell wall damage and mediates apoplastic ROS accumulation in *Arabidopsis* ^56^. Loss-of-function mutants of *PRX71* partially suppress the ROS burst and the associated growth inhibition observed in cell wall–defective mutants, indicating that peroxidase-mediated ROS are essential effectors of the compensatory response. PRX-dependent ROS contribute to oxidative cross-linking of wall polymers and localized lignification, reinforcing tissue mechanics and restricting abnormal expansion. Together, these studies establish ROS as central mediators linking cell wall integrity defects to adaptive growth and developmental responses. In the context of apical hook development, this suggests that cuticle integrity may similarly interact with ROS-mediated mechanical feedback: defects in the cuticle or wall could perturb local tissue mechanics, influencing growth gradients and the maintenance of organ curvature, analogous to the effects observed with cell wall defects and peroxidase-dependent ROS.

While different environmental or developmental cues may activate distinct signaling pathways, they may converge in ROS production, which is known to influence cell wall properties, directional cell expansion, and cortical microtubule dynamics ^69^. One example is the recent work showing that osmotic stress triggers ROS accumulation through a ROP6-dependent pathway activated by the guanine nucleotide exchange factor GEF14, which organizes stimulus-specific ROP6 nanodomains at the plasma membrane ^70^. In this framework, the reduced growth anisotropy and attenuation of antagonistic growth gradients in cuticle-defective apical hooks may be consistent with altered ROS-mediated control of directional growth at the cellular level. Although these links remain speculative, they would suggest that the cuticle can support tissue-scale morphogenesis by modulating the capacity of epidermal cells to generate and maintain coordinated, anisotropic growth patterns.

## Material and methods

### Plant material and growth conditions

All *Arabidopsis thaliana* mutants and transgenic lines were in Columbia-0 (Col-0) ecotype background except *p35S::GFP-MBD* which was in Ws-4 background. *Arabidopsis* lines *dcr-2* (SALK_128228), *cyp77a4-3 pCYP77A4::CYP77A4-GFP*, *lacs1-1* (SALK_127191), *abcg11-7* (+/-) (GABI_590C03), *gpat8* (SALK_035914), *p35S::PIP2A-GFP*, *p35S::GFP-MBD*, *p35S::SUN1-YFP*, *qua2-1*, *gae1-1 gae6-1*, *rbohD* (NASC: N9555), *prx71-1* (SALK_123643) and *35S::PRX71* were described previously ^32,35,36,44,56,71–79^. The *Arabidopsis* lines *bdg1-8* (SM_3_24890) and *cyp77a4-4* (SALK_092049C), were obtained from the publicly available library at NASC ^80^. The selection of *hkb2* and *hkb4* was previously described ^31^. The *cyp77a4-4 p35S::PIP2A-GFP, dcr-2 p35S::PIP2A-GFP, cyp77a4-4 p35S::SUN1-YFP*, *dcr-2 p35S::SUN1-YFP*, *cyp77a4-4 p35S::GFP-MBD, dcr-2 p35S::GFP-MBD*,*dcr-2 prx71-1* and *cyp77a4-4 rbohD* lines were generated by crossing. The homozygotes were screened using fluorescence and PCR genotyping, which was performed by extracting template DNA extracted as previously described ^81^, followed by PCR amplification using primers in Table S2 for insertional mutants. Because *p35S::GFP-MBD* was in the Ws-4 background, F_2_ generation seedlings obtained from the cross between *p35S::GFP-MBD* and *dcr-*2 were screened for fluorescence, and seedlings that genotyped as wild-type for *DCR* were chosen as the control line (Col-0 x Ws-4). *pSAUR16::CDEF1-mCherry*, *pSAUR57::CDEF1-mCherry* and *pDCR::CDEF1-mCherry* in the *p35S::PIP2A-GFP* (Col-0) background, and *dcr-2 pDCR::DCR-mCherry* lines were produced in this study.

Seeds were surface sterilized for two minutes with a solution of 70% ethanol and 1% Tween 20, then washed twice with 95% ethanol and left drying under a sterile hood before sowing on square Petri plates (12x12 cm) of growth medium (GM) composed of half strength Murashige and Skoog (MS) medium (Duchefa Biochemie, Haarlem, The Netherlands), 0.05 % morpholinoethanesulfonic acid (Sigma-Aldrich), 1 % sucrose and 0.8 % agar (Duchefa Biochemie), at pH 5.7. Seeds were placed at 4°C for two to four days and then transferred to 22°C for 16/8 hours of light/darkness per day, or for six hours of light before wrapping in aluminium foil for dark growth. For kinematics of apical hook development, after six hours of light treatment, plates were placed vertically in a dark box equipped with infrared lights (940 nm) (SOLAROX®) and photographed with Raspberry Pi NoIR v2 cameras controlled by Raspberry Pi 3 Model B+ computers. For time-lapse cell length quantification, seedlings were germinated in Petri plates and then transferred to 3D printed square black plates using PETG filaments (size 1.75 mm) containing half strength MS medium. After the transfer, one drop of 1% low melting agar (Invitrogen™ UltraPure™ Low Melting Point Agarose) was added at the bottom of the hypocotyl to maintain the seedlings in place.

### Chemical treatment

A 3 mM stock solution of diphenylene iodonium chloride (DPI; Sigma-Aldrich) was prepared in dimethyl sulfoxide (DMSO) and added to half-strength MS medium at the indicated concentrations during plate preparation. Control medium contained an equivalent volume of DMSO for comparison.

### Sequencing EMS mutants

The *hkb2* and *hkb4* mutants were backcrossed twice with Col-0. Genomic DNA extraction and whole genome sequencing were performed as described previously ^31^.

### Generation of transgenic plants

For all the cloning, DNA amplification by PCR was performed using Phusion High-Fidelity polymerase (Invitrogen), which was then inserted into GreenGate vectors using methods previously described ^82^. The promoter of DCR (2930 bp), SAUR16 (2574 bp) and SAUR57 (3231 bp) were amplified with the following primers: proDCR-F and proDCR-R; proSAUR16-F and proSAUR16-R; proSAUR57-F and proSAUR57-R respectively, and the products were inserted into the pGGA000 vector. The CDEF1 coding sequence (1044 bp) was amplified using CDEF1-F and CDEF1-R primers; the DCR coding sequence (1452 bp) was amplified using DCRFwGG and DCRRevGG, and both the products were inserted in the pGGC000 vector. The primer sequences are listed in Table S2.

The entry vectors were used to generate *pDCR::CDEF1-mCherry*, *pSAUR16::CDEF1-mCherry*, *pSAUR57::CDEF1-mCherry* and *pDCR::DCR-mCherry* binary destination vectors. Stable transformation of *A. thaliana* mediated by *Agrobacterium tumefaciens* was performed by floral dip ^83^. Transformants were selected on hygromycin and homozygous T3 lines were confirmed by the presence of fluorescence in all screened seedlings.

### Toluidine blue staining

Toluidine blue (Sigma 198161) staining was performed as previously described ^33^. Briefly, seedlings were transferred to wells of a 12-well-plate containing 0.1% toluidine blue water solution for two minutes and then washed in distilled water. Seedlings were mounted in water on glass slides and images were acquired with a Zeiss Axioplan microscope.

### Propidium iodide (PI) staining

PI staining (Sigma-Aldrich P4170) was performed to visualize cell walls. Etiolated seedlings (48 hours) were incubated in a freshly prepared PI solution (10 µg/mL ½ MS liquid medium) for 15 mins for Col-0 and 1 minute for *cyp77a4-4* and *dcr-2* at room temperature. Following staining, seedlings were rinsed in ½ MS liquid medium. Seedlings were mounted on glass slides with water and cover slip and Z-stack images were acquired with the Zeiss LSM800 at most 1 um thickness covering the top to the middle of the hypocotyl.

### 3,3’-Diaminobenzidine (DAB) staining

DAB staining was performed as described previously ^84^. The DAB staining solution was prepared by adding 1 mg/ml DAB (Sigma-Aldrich) to sterile water with stirring and 0.2 M hydrochloric acid (HCl) was added drop-wise to bring the pH of the staining solution to 3.0. Subsequently, 25 µl TWEEN^®^ 20 (0.05% v/v) and 2.5 ml 200 mM Na_2_HPO_4_ were added to the solution, with continuous stirring. 2 ml of the DAB staining solution was transferred per well to a 12-well plate, and seedlings were incubated in the solution for 6–7 hours with shaking at 80–100 rpm. 10 mM Na_2_HPO_4_ solution was used as a control. Following the incubation, the staining solution was replaced with clearing solution (ethanol:acetic-acid:glycerol in a 3:1:1 ratio) and the plate was incubated at 65°C for 15 minutes. The bleaching solution was then replaced with fresh bleaching solution and the plate was allowed to stand at room temperature for 30 minutes. The apical hook region was directly visualized for DAB staining under a Leica DMi8 microscope.

### Confocal microscopy

For time-lapse cell length quantification, lines expressing *p35S::PIP2A-GFP* were imaged with a ZEISS LSM800 confocal microscope using the W N-Achroplan 20x/0.5 water-immersion objective, and samples were excited with a 488 nm wavelength laser. Z-stacks were acquired with a step-size of 1 µm, with the stack extending from slightly above the epidermis to the stele. Seedlings were imaged at 2 time-points in an 8-hour interval during the maintenance phase. The first time-point was approximately 24–28 hours after germination, and the second time point was approximately 32–36 hours after germination.

For CYP77A4, DCR and CDEF protein localization, seedlings were mounted on glass slides with liquid half-strength MS medium supplemented with 1% sucrose at pH5.7. Confocal Z-stacks were acquired using a ZEISS LSM800 confocal microscope equipped with a C-Apochromat 40x/1.20 water-dipping objective, and the seedlings were excited with a laser of wavelength 488 nm (*pCYP77A4::CYP77A4-GFP*), 587 nm (*pDCR::DCR-mCherry*) or 561 nm (*pSAUR16::CDEF-mCherry*, *pSAUR57::CDEF-mCherry* and *pDCR::CDEF-mCherry)*.

For lines expressing *p35S::MBD-GFP* and *p35S::SUN1-YFP*, images were acquired with a ZEISS LSM800 confocal microscope using the W Plan-Apochromat 40x/1.0 differential interference contrast (DIC) water-immersion objective, and samples were excited using a 482 nm laser (GFP) or a 508 nm laser (YFP). Z-stack step-size was set to 0.62 µm, with the stacks spanning from slightly above the epidermis to the stele.

For PI staining, fluorescence imaging was performed using a ZEISS LSM800 confocal microscope with excitation at 561 nm and emission detected from 600 nm.

### Image analysis

For quantification of apical hook development, the angle between the cotyledons and the hypocotyl was measured using the angle tool in ImageJ as described previously ^9,10^. For quantification of cell growth patterns using live, time-lapse imaging, the acquired images were processed using MorphoGraphX as described previously ^30^. Microtubule orientation and anisotropy was quantified using the SurfCut2, Segmentation4FTBatch, FibrilTool, Cell_AspectRatio and Angle2Cell plug-ins for ImageJ (v1.54f) ^85^ using the workflow described previously ^49,50^. Briefly, the confocal stacks were processed with SurfCut2 to separately extract the cell boundaries and microtubule signals. The cell boundaries were subsequently segmented using the Segmentation4FTBatch macro. FibrilTool was subsequently used to quantify microtubule organisation. The Cell_Aspect_Ratio macro was then used to define the cell axes, following which the Angle2Cell macro was used to express microtubule organisation in relation to the cell axis. The anisotropy values were then extracted from the output of the final macro. All the macros used in this work have been adapted from the MT_Angle2Ablation_Workflow repository used in Demes and Verger, 2023 ^50^ (https://github.com/VergerLab/MT_Angle2Ablation_Workflow). For the quantification of nuclear shape, acquired Z-stack images were processed using FIJI (ImageJ v1.54f). In brief, Z-projections of acquired confocal images were generated, following which the polygon tool was used to manually demarcate nuclear boundaries. ∼50 nuclei of epidermal cells of the apical hook were outlined per seedling. The ROI (regions of interest) analysis tool built into FIJI was used to calculate the aspect ratio (ratio of primary to secondary axis length) of the best-fit ellipses that followed the manually defined outlines using the “Fit Ellipses” and “Shape Descriptors” measurement options. For statistical testing, a one-way analysis of variance was performed followed by a post-hoc Tukey test for honestly significant difference.

### Transmission electron microscopy

The protocol for transmission electron microscopy was adapted from previously described protocols ^18,86^. With a sharp razor blade, the apical hooks of thirty-hour-old seedlings were separated from the hypocotyl and gently transferred into a 2 mL tube containing 2.5% glutaraldehyde solution in 0.1M phosphate buffer (PB) pH 7.4 fixative for 1 hour at room temperature. After removing the fixative, 1-hour post fixation was performed in 1.5% osmium tetroxide and 1.5% potassium ferrocyanide in water. For sample dehydration, the following series was used: 30% ethanol 15 minutes, 50% ethanol 15 minutes, 70% ethanol 15 minutes, and 100% ethanol 15 minutes twice. Spurr resin (TAAB Laboratories, Aldermaston, England) infiltration was then performed with 33% resin in ethanol for 2 hours, 66% resin in ethanol for 2 hours, and finally 100% for 8 hours twice. Single hooks were aspirated with a plastic Pasteur pipette and placed at the bottom of an embedding capsule (TAAB Laboratories, Aldermaston, England) and 100% fresh spurr resin was added on top of the sample, filling the capsule. Samples were polymerized at 60 °C for 24 hours. 70 nm ultrathin sections were cut and then picked up on copper grids and examined with a Talos L120C transmission electron microscope (FEI, Eindhoven, The Netherlands) operating at 120 kV. Micrographs were acquired with a Ceta 16M CCD camera (FEI, Eindhoven, The Netherlands) using Velox ver 2.14.2.40.

### Atomic force microscopy (AFM)

Samples were mounted on a small petri dish with a thin layer of FLEXBAR Reprorubber® Thin Pour Material (Flexbar Machine Corporation) to ensure sample immobilization without covering the sample surface. Once the layer was solidified, the samples were immersed in water. The AFM experiments were performed using a NanoWizard® 4 XP BioScience AFM (Bruker) equipped with HybridStageTM controlled by JPK SPMControl Software version 7. The petri dish was placed on the AFM stage and the bottom part of the hypocotyl was localized using the brightfield optical system and camera of the microscope. Direct AFM measurements in the apical hook region were technically challenging as the cotyledons prevented the samples from lying completely flat and interfered with cantilever approach during acquisition. A low-resolution Quantitative Imaging™ (QI) map with an image size of 90 × 180 µm and a resolution of 45 × 90 pixels (2 µm per pixel) was acquired on the hypocotyls to identify the cells for Young’s modulus measurements. Once the cells were visible on the QI map, the mode of the AFM was switched to Contact Force Spectroscopy and 30-40 measurement points were defined per cell, following which the force spectra measurements were recorded. Two regions of interest were acquired using QI mode for each hypocotyl per genotype per biological replicate. Three biological replicates were performed. The experiments were conducted in water at room temperature using biosphere B500-NCH cantilevers, which have a spring constant of 40 N/m and a tip diameter of 0.5 µm, with a sphere, Diamond-Like-Carbon tip. The cantilever calibration was done using a pre-calibrated method in JPK SPMControl Software version 7. The QI maps were acquired with the following settings: setpoint 500 nN, pixel time 65 ms, Z length 4.5 µm, speed 173.08 µm/s and all Contact Mode Force Spectra measurements were recorded with the following settings: setpoint 600 nN, Z Length 2 µm, and the Z Speed 1 µm/s. Contact Mode Force Spectra measurements were analyzed using the JPK Data Processing Software. To extract the apparent Young’s modulus, the following functions were applied on the measurements: calibration, baseline subtraction, contact point and vertical tip position. The Hert/Sneddon model was first fitted to the entire force-depth curve and then refitted within the ∞ -100 indentation range using following settings: spherical tip shape, 500 nm tip radius and 0.5 Poisson’s ratio. Once the apparent Young’s modulus values were obtained, one mean value was calculated for each cell.

### Finite element model (FEM)

To investigate the biomechanical relationship between local cell expansion patterns and apical hook morphogenesis, a two-dimensional finite element model (FEM) of the apical hook was developed, that simulates morphoelastic growth dynamics. The model was implemented in BVPy, a Python library based on FEniCS, which enables the numerical solution of partial differential equations using FEM. The model geometry was reconstructed from MorphoGraphX and projected on a 2D surface, segmented into cellular and extracellular domains. Each cellular domain corresponded to one biological cell, while the extracellular space represented the shared anticlinal walls of neighboring cells in a 2D projection. Growth was described by assigning a strain rate tensor to each cellular domain, derived from experimentally measured growth patterns. This allowed each cell within the mesh to express a unique, anisotropic growth scheme consistent with observed expansion behavior. The extracellular domain exhibited strain-dependent growth, coupling deformation with local mechanical stress to represent the passive mechanical response of the cell wall matrix. To account for spatial mechanical heterogeneity, two regional variations in Young’s modulus were defined. These differences reflected the contribution of anticlinal walls to the overall stiffness of the 2D tissue representation. Poisson’s ratio (0.4) was assumed to be constant across the tissue domain. Growth and deformation were coupled using a morphoelastic formulation based on the St. Venant–Kirchhoff hyperelastic material model. Dirichlet boundary conditions were applied by fixing the basal region of the hook to prevent translation or rotation, allowing deformation to occur only in the upper tissue regions. The FEM solver computed the equilibrium displacement field at each iteration with a time resolution of 2.4 min over 8 hours.

### Statistical analysis

Statistical analyses were performed in R (v4.5.1). The effect of genotypes, domains, and their interaction on observations were tested using two-way linear model fitted with the “*stats*” package. Adjusted means for all combinations were computed using “*emmeans*” package. Multiple comparisons among these combinations were summarized using a compact letter display generated with the “*multcomp*” package, with Šidák correction applied.

## Acknowledgements

We acknowledge the Knut and Alice Wallenberg Foundation for essential financing of Umeå Plant Science Centre (UPSC) via Wallenberg Initiatives in Forest Research (WIFORCE). We acknowledge the UPSC Microscopy Facility for access to imaging equipment and the UPSC Bioinformatics Facility (https://bioinfomatics.upsc.se) for technical support with regards to computational resources used for the FEM and the Umeå Centre for Electron Microscopy (UCEM) for assistance with TEM imaging. We thank Olivier Ali for feedback on the modeling strategy. This work was financially supported by the Kempe Foundation (S.Ra., INUPRAG to F.J.; JCK-2130 to A.H.; JCK-1732 to S.L.; JCK-1912.2; A.A.), the Swedish Research Council (VR 2020-03420; VR 2024-04028; VR 2013-4632; F.J., S.Ro., H.R., VR, 202003974; A.A.), Stiftelsen Olle Engkvist Byggmästare (S.Ra; 185 595), the Knut and Alice Wallenberg Foundation (S.L.; FATE, 2022.0029, S.Ra., L.L.D.), VINNOVA (Verket för Innovationssystem) (S.L., S.Ra., H.R., S.Ro.) and the European Research Council (ERC-2024-SyG STARMORPH 101166880 to S.M.D. S.Ro. and J.K.-V.). This work has been funded by Ministerio de Ciencia Innovación y Universidades of Spain (PID2024-155159NB-I00, CNS2023-143915; KW), and Severo Ochoa (SO) Program for Centers of Excellence in R&D from the Agencia Estatal de Investigación of Spain [grant CEX2020-000999-S (2022 to 2025) to the CBGP]. This work was also supported by funding from the Deutsche Forschungsgemeinschaft (DFG) (DFG; 499026372 to J.K.-V. and CIBSS – EXC-2189; 390939984 to J.K.-V).

## Author contributions

S.Ra., H.R. and S.Ro. designed the research. S.Ra., H.R., L.L.D., Ö.E., S.M.D and F.J. performed the experiments under the guidance of S.Ro., S.V. and J.K.V. A.A. and S.L. performed preliminary data acquisition under the guidance of S.V. and S.Ro. A.H. performed the computer simulations assisted by H.R. under the guidance of M.P., K.W. and S.Ro. All authors analyzed the data. S.Ra., H.R. and S.Ro. wrote the article. All authors edited the final manuscript.

## Competing interests

The authors declare no competing financial interests.

## Additional information

**Extended data** for this paper is available online.

**Supplementary information:** The online version contains supplementary material.

**Correspondence and requests for materials** should be addressed to Stéphanie Robert.

---

**Supplementary Figure 1.**
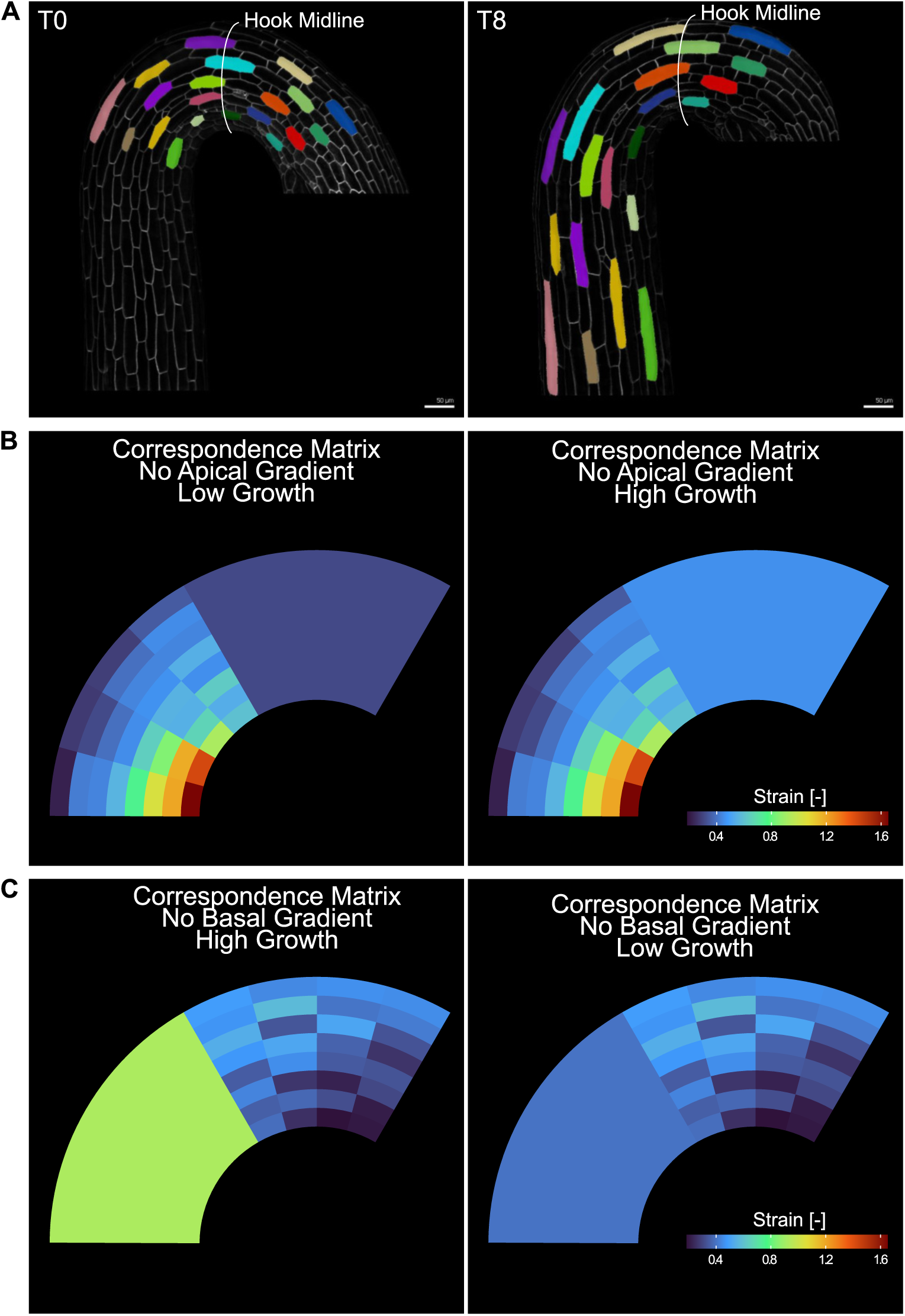
Experimental and modeling frameworks underlying apical hook maintenance and growth parameterization. (A) Representative live confocal micrographs of dark-grown *Arabidopsis* apical hooks expressing *p35S::PIP2A-GFP* at approximately 24 h after germination (T0) and 8 h later (T8), illustrating hook maintenance during this developmental window. Selected cells are highlighted to visualize cell flow over time. Scale bars represent 50 µm. (B) Correspondence matrices used to conduct simulations lacking an apical growth gradient, shown for two growth scenarios: low and high growth. (C) Correspondence matrices used to conduct simulations lacking a basal growth gradient, shown for two growth scenarios: low and high growth. Panels (B) and (C) illustrate the growth properties used in Fig. 1F.

**Supplementary Figure 2.**
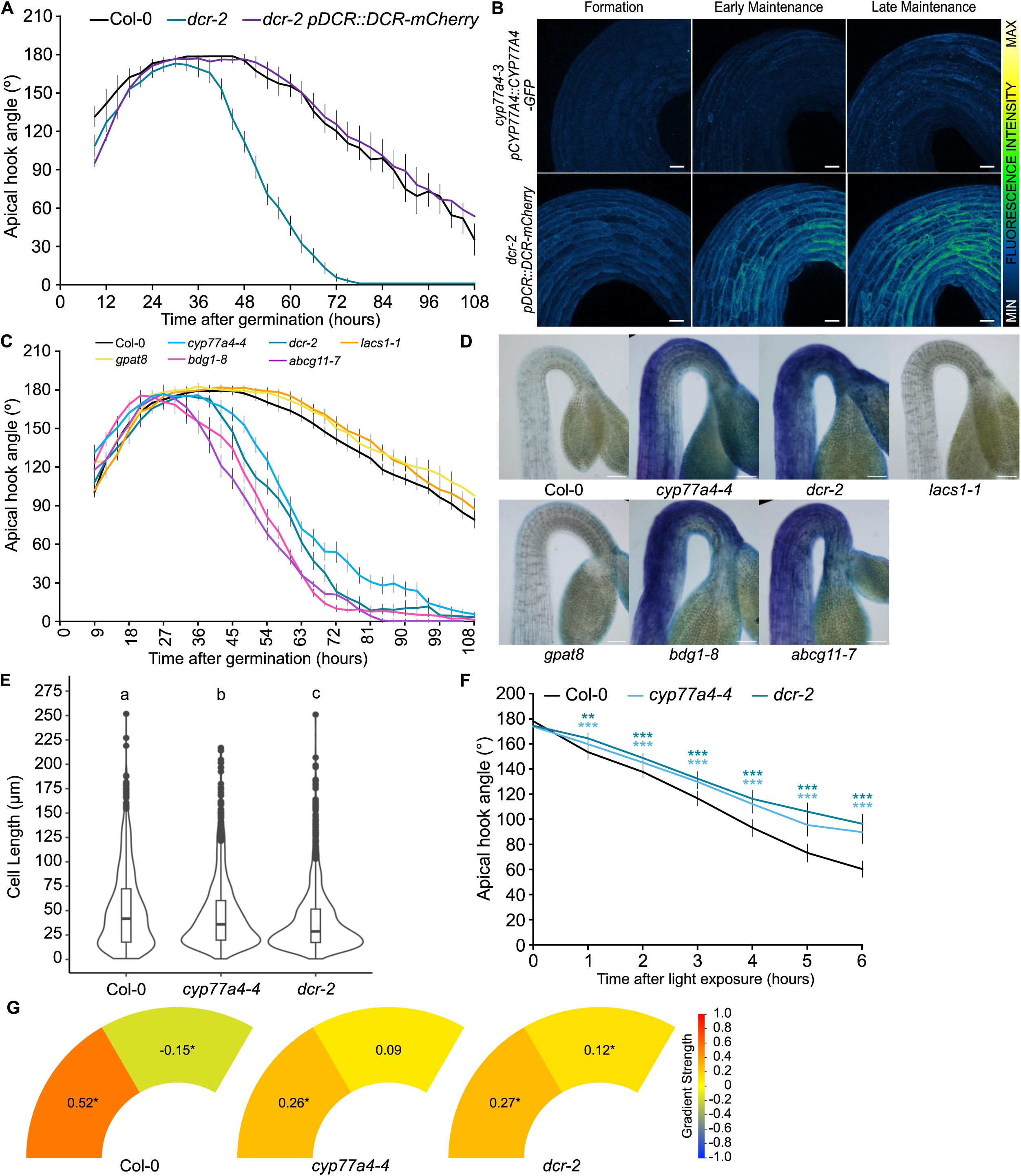
Cuticle integrity is essential for apical hook development. (A) Kinematics of apical hook angle in dark-grown Col-0 wild-type, *dcr-2* and *dcr-2 pDCR::DCR-mCherry* as measured every 3 h, with 0 h as the time point of germination. The means of 7–15 seedlings per line are shown; error bars represent standard error of the mean. (B) Representative live confocal micrographs of dark-grown apical hooks of *cyp77a4-3 pCYP77A4::CYP77A4-GFP* and *dcr-2 pDCR::DCR-mCherry* in the formation, early maintenance and late maintenance phases (approximately 24, 36 and 48 h after germination, respectively). Scale bars represent 20 µm. Increasing fluorescence intensity is color-coded according to the scale shown. (C) Kinematics of apical hook angle in dark-grown Col-0 wild-type, *cyp77a4-4*, *dcr-2*, *lacs1-1*, *gpat8*, *bdg1-8* and *abcg11-7*, as measured every 3 h, with 0 h as the time point of germination. The means of 11 to 70 seedlings per line are shown; error bars represent standard error of the mean. (D) Representative light micrographs of dark-grown upper hypocotyls and apical hooks of the lines shown in (C) in the maintenance phase (48 hours post-germination) after toluidine blue staining. Scale bars represent 50 µm. (E) Violin plots of cell length for each domain shown in apical hooks expressing stained with PI in Col-0 wild-type, *cyp77a4-4* and *dcr-2* backgrounds at approximately 48 h after germination. 900–1600 cells per genotype were measured over at least three biological replicates. Different letters indicate statistically significant differences based on one-way ANOVA followed by Tukey’s *post-hoc* tests (α = 0.05). (F) Kinematics of apical hook angle in dark-grown Col-0 wild-type, *cyp77a4-4* and *dcr-2* upon treatment with light, as measured every 1 h, with 0 h as the time point of light exposure. Seedlings were light-treated at 24 h post-germination. The means of 20 seedlings per line are shown; error bars represent standard error of the mean. Asterisks indicate significant difference of *cyp77a4-4* (light blue) or *dcr-2* (dark blue) to Col-0 according to the Student’s t-test (\*\**P* < 0.01, \*\*\**P* < 0.001). (G) Schematics representing apical and basal elongation gradient strength in Col-0 wild-type, *cyp77a4-4* and *dcr-2*, expressed as a percentage ratio of average longitudinal strains in the inner domain to the outer domain. A positive value indicates that the inner domain has a higher strain than the outer domain and *vice-versa*. Asterisks represent significant differences in average longitudinal strain in comparisons between inner and outer domains. A linear model with interaction effects between genotypes and domains was used for a pairwise comparison followed by a Dunn-Šidák correction (α = 0.05).

**Supplementary Figure 3.**
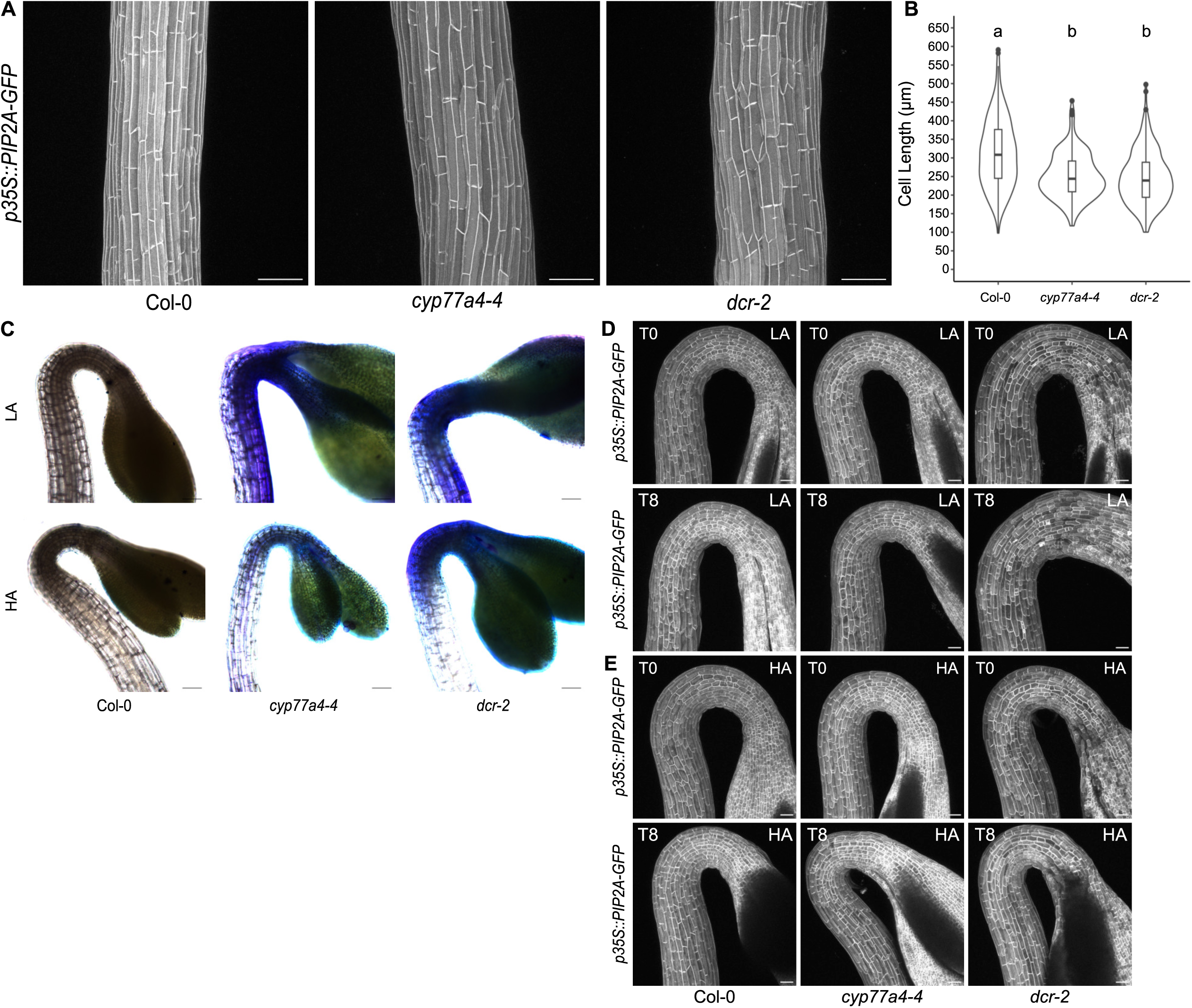
Mechanical cues link cuticle integrity to apical hook development. (A) Representative live confocal micrographs of basal regions of dark-grown hypocotyls of *p35S::PIP2A-GFP* in Col-0 wild-type, *cyp77a4-4* and *dcr-2* backgrounds. Scale bars represent 100 µm. (B) Violin plots of epidermal cell length in basal regions of hypocotyls expressing *p35S::PIP2A-GFP* in Col-0 wild-type, *cyp77a4-4* and *dcr-2* backgrounds, see representative images in (A). 10–20 cells were measured for each of ≥15 seedlings per line across 3 biological replicates. Different letters indicate statistically significant differences based on one-way ANOVA followed by Tukey’s *post-hoc* tests (α = 0.05). (C) Representative light micrographs of dark-grown upper hypocotyls and apical hooks of Col-0, *cyp77a4-4* and *dcr-2* grown on medium with low-agar (LA, 0.8% w/v agar) and high-agar (HA, 2.5% w/v agar) in the maintenance phase (48 hours post-germination) after toluidine blue staining. Scale bars represent 100 µm. (D-E) Representative live confocal micrographs of dark-grown apical hooks expressing the plasma membrane marker *p35S::PIP2A-GFP* in Col-0 wild-type, *cyp77a4-4* and *dcr-2* backgrounds grown on LA (D) and HA (E) at approximately 24 h after germination (T0) and 8 h later (T8). Scale bars represent 50 µm.

**Supplementary Figure 4.**
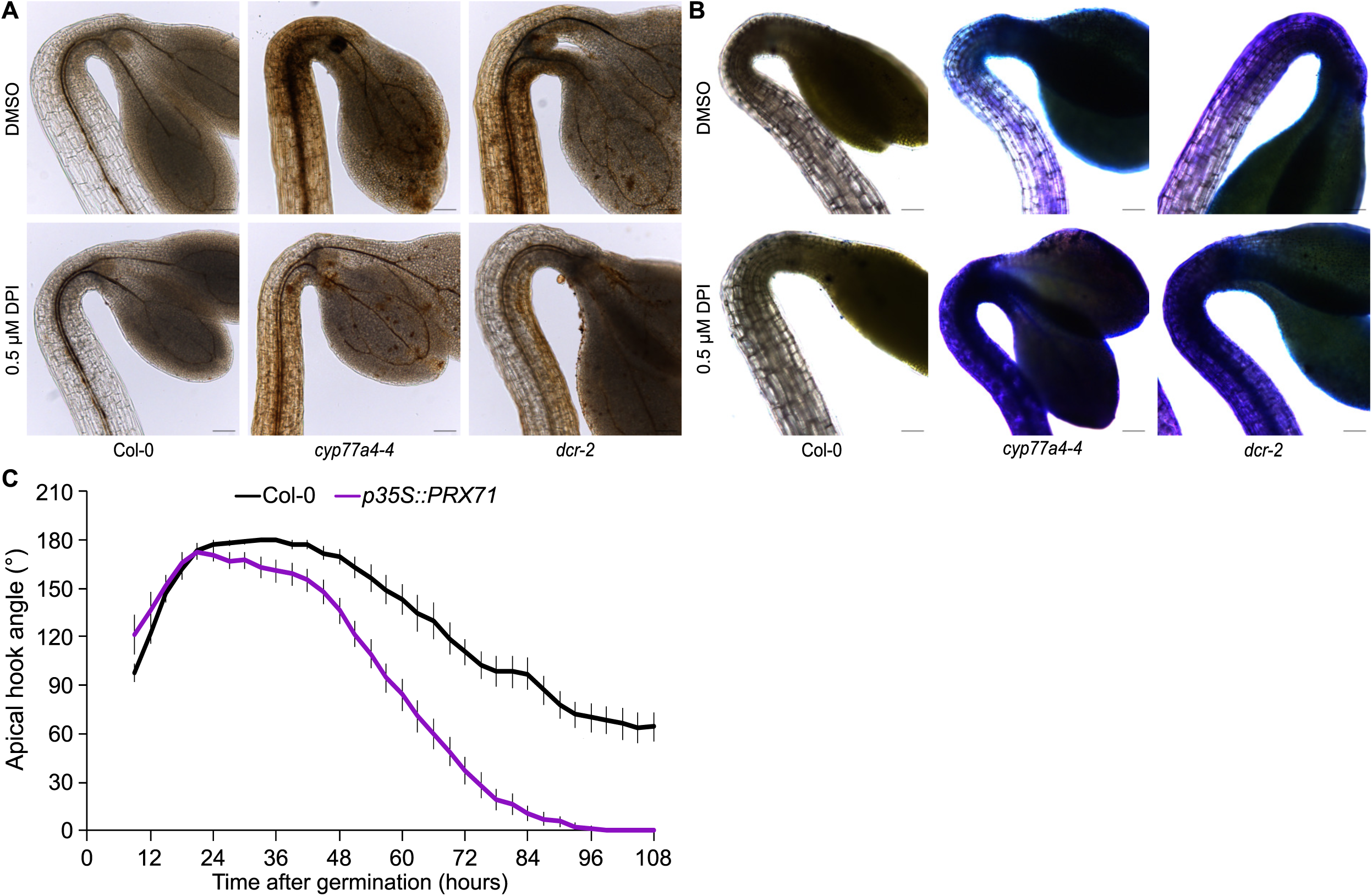
Reactive oxygen species (ROS) link biomechanical cues with cuticle permeability during apical hook maintenance. (A-B) Representative light micrographs of dark-grown upper hypocotyls and apical hooks of Col-0, *cyp77a4-4* and *dcr-2* grown on medium supplemented with 0.5 µM DPI in the maintenance phase (48 hours post-germination) after DAB staining (A) or toluidine blue staining (B). Scale bars represent 100 µm. 5 to 8 seedlings were analyzed for DAB staining over 3 biological replicates. (C) Kinematics of apical hook angle in dark-grown Col-0 wild-type and *p35S::PRX71* as measured every 3 h, with 0 h as the time point of germination. The means of 12–19 seedlings per line are shown; error bars represent standard error of the mean.

## Notes

### Competing Interest Statement

The authors have declared no competing interest.

### Summary of Updates

The acknowledgements section had to be updated to reflect recent changes in the author list. The previous version contained grants and funding sources associated with the author that has been dropped from the manuscript, which have now been appropriately amended.

## References

1. Geitmann, A., and Ortega, J.K.E. (2009). Mechanics and modeling of plant cell growth. Trends in Plant Science 14, 467–478. 10.1016/j.tplants.2009.07.006.

2. Bidhendi, A.J., and Geitmann, A. (2016). Relating the mechanics of the primary plant cell wall to morphogenesis. J Exp Bot 67, 449–461. 10.1093/jxb/erv535.

3. Cosgrove, D.J. (2016). Plant cell wall extensibility: connecting plant cell growth with cell wall structure, mechanics, and the action of wall-modifying enzymes. J Exp Bot 67, 463–476. 10.1093/jxb/erv511.

4. Cosgrove, D.J. (2024). Plant Cell Wall Loosening by Expansins. Annual Review of Cell and Developmental Biology 40, 329–352. 10.1146/annurev-cellbio-111822-115334.

5. Delmer, D., Dixon, R.A., Keegstra, K., and Mohnen, D. (2024). The plant cell wall—dynamic, strong, and adaptable—is a natural shapeshifter. Plant Cell 36, 1257–1311. 10.1093/plcell/koad325.

6. Majda, M., Grones, P., Sintorn, I.-M., Vain, T., Milani, P., Krupinski, P., Zagórska-Marek, B., Viotti, C., Jönsson, H., Mellerowicz, E.J., et al. (2017). Mechanochemical Polarization of Contiguous Cell Walls Shapes Plant Pavement Cells. Developmental Cell 43, 290–304.e4. 10.1016/j.devcel.2017.10.017.

7. Majda, M., and Robert, S. (2018). The Role of Auxin in Cell Wall Expansion. International Journal of Molecular Sciences 19. 10.3390/ijms19040951.

8. Raz, V., and Ecker, J.R. (1999). Regulation of differential growth in the apical hook of Arabidopsis. Development 126, 3661–3668. 10.1242/dev.126.16.3661.

9. Žádníková, P., Petrášek, J., Marhavý, P., Raz, V., Vandenbussche, F., Ding, Z., Schwarzerová, K., Morita, M.T., Tasaka, M., Hejátko, J., et al. (2010). Role of PIN-mediated auxin efflux in apical hook development of Arabidopsis thaliana. Development 137, 607–617. 10.1242/dev.041277.

10. Ma, Q., Liu, S., Doyle, S.M., Raggi, S., Pařízková, B., Barange, D.K., Ratnakaram, H., Wilkinson, E.G., Crespo Garcia, I., Bygdell, J., et al. (2025). RACK1A positively regulates opening of the apical hook in Arabidopsis thaliana via suppression of its auxin response gradient. Proceedings of the National Academy of Sciences 122, e2407224122. 10.1073/pnas.2407224122.

11. Abbas, M., Alabadí, D., and Blázquez, M.A. (2013). Differential growth at the apical hook: all roads lead to auxin. Front Plant Sci 4, 441. 10.3389/fpls.2013.00441.

12. Wang, Y., Peng, Y., and Guo, H. (2023). To curve for survival: Apical hook development. Journal of Integrative Plant Biology 65, 324–342. 10.1111/jipb.13441.

13. Willoughby, A.C., and Strader, L.C. (2024). Apical hook opening of plant seedlings: Unfolding the role of auxin and the cell wall. Developmental Cell 59, 3194–3196. 10.1016/j.devcel.2024.11.018.

14. Walia, A., Carter, R., Wightman, R., Meyerowitz, E.M., Jönsson, H., and Jones, A.M. (2024). Differential growth is an emergent property of mechanochemical feedback mechanisms in curved plant organs. Developmental Cell 59, 3245–3258.e3. 10.1016/j.devcel.2024.09.021.

15. Béziat, C., and Kleine-Vehn, J. (2018). The Road to Auxin-Dependent Growth Repression and Promotion in Apical Hooks. Current Biology 28, R519–R525. 10.1016/j.cub.2018.01.069.

16. Sassi, M., Ali, O., Boudon, F., Cloarec, G., Abad, U., Cellier, C., Chen, X., Gilles, B., Milani, P., Friml, J., et al. (2014). An Auxin-Mediated Shift toward Growth Isotropy Promotes Organ Formation at the Shoot Meristem in *Arabidopsis*. Current Biology 24, 2335–2342. 10.1016/j.cub.2014.08.036.

17. Yeats, T.H., and Rose, J.K.C. (2013). The Formation and Function of Plant Cuticles. Plant Physiol 163, 5–20. 10.1104/pp.113.222737.

18. Berhin, A., de Bellis, D., Franke, R.B., Buono, R.A., Nowack, M.K., and Nawrath, C. (2019). The Root Cap Cuticle: A Cell Wall Structure for Seedling Establishment and Lateral Root Formation. Cell 176, 1367–1378.e8. 10.1016/j.cell.2019.01.005.

19. Renault, H., Alber, A., Horst, N.A., Basilio Lopes, A., Fich, E.A., Kriegshauser, L., Wiedemann, G., Ullmann, P., Herrgott, L., Erhardt, M., et al. (2017). A phenol-enriched cuticle is ancestral to lignin evolution in land plants. Nat Commun 8, 14713. 10.1038/ncomms14713.

20. Ingram, G., and Nawrath, C. (2017). The roles of the cuticle in plant development: organ adhesions and beyond. J Exp Bot 68, 5307–5321. 10.1093/jxb/erx313.

21. Foreman, J., Demidchik, V., Bothwell, J.H.F., Mylona, P., Miedema, H., Torres, M.A., Linstead, P., Costa, S., Brownlee, C., Jones, J.D.G., et al. (2003). Reactive oxygen species produced by NADPH oxidase regulate plant cell growth. Nature 422, 442–446. 10.1038/nature01485.

22. Monshausen, G.B., Bibikova, T.N., Messerli, M.A., Shi, C., and Gilroy, S. (2007). Oscillations in extracellular pH and reactive oxygen species modulate tip growth of Arabidopsis root hairs. Proceedings of the National Academy of Sciences 104, 20996–21001. 10.1073/pnas.0708586104.

23. Mangano, S., Denita-Juarez, S.P., Choi, H.-S., Marzol, E., Hwang, Y., Ranocha, P., Velasquez, S.M., Borassi, C., Barberini, M.L., Aptekmann, A.A., et al. (2017). Molecular link between auxin and ROS-mediated polar growth. Proceedings of the National Academy of Sciences 114, 5289–5294. 10.1073/pnas.1701536114.

24. Waszczak, C., Carmody, M., and Kangasjärvi, J. (2018). Reactive Oxygen Species in Plant Signaling. Annual Review of Plant Biology 69, 209–236. 10.1146/annurev-arplant-042817-040322.

25. Survila, M., Davidsson, P.R., Pennanen, V., Kariola, T., Broberg, M., Sipari, N., Heino, P., and Palva, E.T. (2016). Peroxidase-Generated Apoplastic ROS Impair Cuticle Integrity and Contribute to DAMP-Elicited Defenses. Front. Plant Sci. 7. 10.3389/fpls.2016.01945.

26. L’Haridon, F., Besson-Bard, A., Binda, M., Serrano, M., Abou-Mansour, E., Balet, F., Schoonbeek, H.-J., Hess, S., Mir, R., Léon, J., et al. (2011). A Permeable Cuticle Is Associated with the Release of Reactive Oxygen Species and Induction of Innate Immunity. PLoS Pathog 7, e1002148. 10.1371/journal.ppat.1002148.

27. Rubinstein, B. (1971). The Role of Various Regions of the Bean Hypocotyl on Red Light-induced Hook Opening. Plant Physiol. 48, 183–186. 10.1104/pp.48.2.183.

28. Silk, W.K., and Erickson, R.O. (1978). KINEMATICS OF HYPOCOTYL CURVATURE. American J of Botany 65, 310–319. 10.1002/j.1537-2197.1978.tb06072.x.

29. Wang, Y., and Guo, H. (2019). On hormonal regulation of the dynamic apical hook development. New Phytol 222, 1230–1234. 10.1111/nph.15626.

30. Barbier De Reuille, P., Routier-Kierzkowska, A.-L., Kierzkowski, D., Bassel, G.W., Schüpbach, T., Tauriello, G., Bajpai, N., Strauss, S., Weber, A., Kiss, A., et al. (2015). MorphoGraphX: A platform for quantifying morphogenesis in 4D. eLife 4, e05864. 10.7554/eLife.05864.

31. Vain, T., Raggi, S., Ferro, N., Barange, D.K., Kieffer, M., Ma, Q., Doyle, S.M., Thelander, M., Pařízková, B., Novák, O., et al. (2019). Selective auxin agonists induce specific AUX/IAA protein degradation to modulate plant development. Proc. Natl. Acad. Sci. U.S.A. 116, 6463–6472. 10.1073/pnas.1809037116.

32. Kawade, K., Li, Y., Koga, H., Sawada, Y., Okamoto, M., Kuwahara, A., Tsukaya, H., and Hirai, M.Y. (2018). The cytochrome P450 CYP77A4 is involved in auxin-mediated patterning of the Arabidopsis thaliana embryo. Development 145, dev168369. 10.1242/dev.168369.

33. Tanaka, T., Tanaka, H., Machida, C., Watanabe, M., and Machida, Y. (2004). A new method for rapid visualization of defects in leaf cuticle reveals five intrinsic patterns of surface defects in *Arabidopsis*. The Plant Journal 37, 139–146. 10.1046/j.1365-313X.2003.01946.x.

34. Takahashi, K., Shimada, T., Kondo, M., Tamai, A., Mori, M., Nishimura, M., and Hara-Nishimura, I. (2010). Ectopic Expression of an Esterase, Which is a Candidate for the Unidentified Plant Cutinase, Causes Cuticular Defects in Arabidopsis thaliana. Plant and Cell Physiology 51, 123–131. 10.1093/pcp/pcp173.

35. Panikashvili, D., Shi, J.X., Schreiber, L., and Aharoni, A. (2009). The Arabidopsis *DCR* Encoding a Soluble BAHD Acyltransferase Is Required for Cutin Polyester Formation and Seed Hydration Properties. Plant Physiology 151, 1773–1789. 10.1104/pp.109.143388.

36. Wu, L., Zhou, Z.-Y., Zhang, C.-G., Chai, J., Zhou, Q., Wang, L., Hirnerová, E., Mrvková, M., Novák, O., and Guo, G.-Q. (2015). Functional roles of three cutin biosynthetic acyltransferases in cytokinin responses and skotomorphogenesis. PLoS One 10, e0121943. 10.1371/journal.pone.0121943.

37. Wang, J., Sun, N., Zhang, F., Yu, R., Chen, H., Deng, X.W., and Wei, N. (2020). SAUR17 and SAUR50 Differentially Regulate PP2C-D1 during Apical Hook Development and Cotyledon Opening in Arabidopsis. Plant Cell 32, 3792–3811. 10.1105/tpc.20.00283.

38. Rabe, C., and Kutschera, U. (1999). Rapid Light-Induced Enhancement of Sucrose Catabolism in the Apical Hook of Sunflower Hypocotyls. Journal of Plant Physiology 155, 538–542. 10.1016/S0176-1617(99)80051-0.

39. Cosgrove, D.J. (2022). Building an extensible cell wall. Plant Physiology 189, 1246–1277. 10.1093/plphys/kiac184.

40. Belteton, S.A., Li, W., Yanagisawa, M., Hatam, F.A., Quinn, M.I., Szymanski, M.K., Marley, M.W., Turner, J.A., and Szymanski, D.B. (2021). Real-time conversion of tissue-scale mechanical forces into an interdigitated growth pattern. Nat. Plants 7, 826–841. 10.1038/s41477-021-00931-z.

41. Bidhendi, A.J., Altartouri, B., Gosselin, F.P., and Geitmann, A. (2019). Mechanical Stress Initiates and Sustains the Morphogenesis of Wavy Leaf Epidermal Cells. Cell Reports 28, 1237–1250.e6. 10.1016/j.celrep.2019.07.006.

42. Chen, S., Burda, I., Jani, P., Pendrak, B., Silberstein, M.N., and Roeder, A.H.K. (2025). Fibrous network nature of plant cell walls enables tunable mechanics for development. Nat Commun 16, 7565. 10.1038/s41467-025-62844-1.

43. Coen, E., and Cosgrove, D.J. (2023). The mechanics of plant morphogenesis. Science 379. 10.1126/science.ade8055.

44. Fal, K., Korsbo, N., Alonso-Serra, J., Teles, J., Liu, M., Refahi, Y., Chabouté, M.-E., Jönsson, H., and Hamant, O. (2021). Tissue folding at the organ–meristem boundary results in nuclear compression and chromatin compaction. Proc Natl Acad Sci U S A 118, e2017859118. 10.1073/pnas.2017859118.

45. Verger, S., Long, Y., Boudaoud, A., and Hamant, O. (2018). A tension-adhesion feedback loop in plant epidermis. eLife 7, e34460. 10.7554/eLife.34460.

46. Hamant, O., Heisler, M.G., Jönsson, H., Krupinski, P., Uyttewaal, M., Bokov, P., Corson, F., Sahlin, P., Boudaoud, A., Meyerowitz, E.M., et al. (2008). Developmental Patterning by Mechanical Signals in *Arabidopsis*. Science 322, 1650–1655. 10.1126/science.1165594.

47. Sampathkumar, A., Krupinski, P., Wightman, R., Milani, P., Berquand, A., Boudaoud, A., Hamant, O., Jönsson, H., and Meyerowitz, E.M. (2014). Subcellular and supracellular mechanical stress prescribes cytoskeleton behavior in Arabidopsis cotyledon pavement cells. eLife 3, e01967. 10.7554/eLife.01967.

48. Robinson, S., and Kuhlemeier, C. (2018). Global Compression Reorients Cortical Microtubules in Arabidopsis Hypocotyl Epidermis and Promotes Growth. Current Biology 28, 1794–1802.e2. 10.1016/j.cub.2018.04.028.

49. Boudaoud, A., Burian, A., Borowska-Wykręt, D., Uyttewaal, M., Wrzalik, R., Kwiatkowska, D., and Hamant, O. (2014). FibrilTool, an ImageJ plug-in to quantify fibrillar structures in raw microscopy images. Nat Protoc 9, 457–463. 10.1038/nprot.2014.024.

50. Demes, E., and Verger, S. (2023). High-throughput characterization of cortical microtubule arrays response to anisotropic tensile stress. BMC Biol 21, 154. 10.1186/s12915-023-01654-7.

51. Lorrai, R., Erguvan, Ö., Raggi, S., Jonsson, K., Široká, J., Tarkowská, D., Novák, O., Griffiths, J., Jones, A.M., Verger, S., et al. (2024). Cell wall integrity modulates HOOKLESS1 and PHYTOCHROME INTERACTING FACTOR4 expression controlling apical hook formation. Plant Physiol 196, 1562–1578. 10.1093/plphys/kiae370.

52. Lorrai, R., Francocci, F., Gully, K., Martens, H.J., De Lorenzo, G., Nawrath, C., and Ferrari, S. (2021). Impaired Cuticle Functionality and Robust Resistance to Botrytis cinerea in Arabidopsis thaliana Plants With Altered Homogalacturonan Integrity Are Dependent on the Class III Peroxidase AtPRX71. Front. Plant Sci. 12, 696955. 10.3389/fpls.2021.696955.

53. Chen, J., Liao, Y., and Wang, Y. (2025). HyPer7 imaging Unveils the role of hydrogen peroxide in Arabidopsis apical hook development. Biochemical and Biophysical Research Communications 774, 152106. 10.1016/j.bbrc.2025.152106.

54. Thordal-Christensen, H., Zhang, Z., Wei, Y., and Collinge, D.B. (1997). Subcellular localization of H_2_ O_2_ in plants. H_2_ O_2_ accumulation in papillae and hypersensitive response during the barley—powdery mildew interaction. The Plant Journal 11, 1187–1194. 10.1046/j.1365-313X.1997.11061187.x.

55. Li, Y., and Trush, M.A. (1998). Diphenyleneiodonium, an NAD(P)H Oxidase Inhibitor, also Potently Inhibits Mitochondrial Reactive Oxygen Species Production. Biochemical and Biophysical Research Communications 253, 295–299. 10.1006/bbrc.1998.9729.

56. Raggi, S., Ferrarini, A., Delledonne, M., Dunand, C., Ranocha, P., De Lorenzo, G., Cervone, F., and Ferrari, S. (2015). The Arabidopsis Class III Peroxidase AtPRX71 Negatively Regulates Growth under Physiological Conditions and in Response to Cell Wall Damage. Plant Physiol 169, 2513–2525. 10.1104/pp.15.01464.

57. Žádníková, P., Wabnik, K., Abuzeineh, A., Gallemi, M., Van Der Straeten, D., Smith, R.S., Inzé, D., Friml, J., Prusinkiewicz, P., and Benková, E. (2016). A Model of Differential Growth-Guided Apical Hook Formation in Plants. Plant Cell 28, 2464–2477. 10.1105/tpc.15.00569.

58. Van De Poel, B., Smet, D., and Van Der Straeten, D. (2015). Ethylene and Hormonal Cross Talk in Vegetative Growth and Development. Plant Physiol. 169, 61–72. 10.1104/pp.15.00724.

59. Narukawa, H., Yokoyama, R., and Nishitani, K. (2016). Possible pathways linking ploidy level to cell elongation and cuticular function in hypocotyls of dark-grown Arabidopsis seedlings. Plant Signaling & Behavior 11, e1118597. 10.1080/15592324.2015.1118597.

60. Moyroud, E., Airoldi, C.A., Ferria, J., Giorio, C., Steimer, S.S., Rudall, P.J., Prychid, C.J., Halliwell, S., Walker, J.F., Robinson, S., et al. (2022). Cuticle chemistry drives the development of diffraction gratings on the surface of Hibiscus trionum petals. Current Biology 32, 5323–5334.e6. 10.1016/j.cub.2022.10.065.

61. Riglet, L., Gatti, S., and Moyroud, E. (2021). Sculpting the surface: Structural patterning of plant epidermis. iScience 24, 103346. 10.1016/j.isci.2021.103346.

62. Domínguez, E., Heredia-Guerrero, J.A., and Heredia, A. (2011). The biophysical design of plant cuticles: an overview. New Phytologist 189, 938–949. 10.1111/j.1469-8137.2010.03553.x.

63. Knoche, M., and Lang, A. (2017). Ongoing Growth Challenges Fruit Skin Integrity. Critical Reviews in Plant Sciences 36, 190–215. 10.1080/07352689.2017.1369333.

64. Jiang, F., Lopez, A., Jeon, S., De Freitas, S.T., Yu, Q., Wu, Z., Labavitch, J.M., Tian, S., Powell, A.L.T., and Mitcham, E. (2019). Disassembly of the fruit cell wall by the ripening-associated polygalacturonase and expansin influences tomato cracking. Hortic Res 6, 17. 10.1038/s41438-018-0105-3.

65. Reynoud, N., Geneix, N., D’Orlando, A., Petit, J., Mathurin, J., Deniset-Besseau, A., Marion, D., Rothan, C., Lahaye, M., and Bakan, B. (2023). Cuticle architecture and mechanical properties: a functional relationship delineated through correlated multimodal imaging. New Phytologist 238, 2033–2046. 10.1111/nph.18862.

66. Hématy, K., Sado, P.-E., Van Tuinen, A., Rochange, S., Desnos, T., Balzergue, S., Pelletier, S., Renou, J.-P., and Höfte, H. (2007). A Receptor-like Kinase Mediates the Response of Arabidopsis Cells to the Inhibition of Cellulose Synthesis. Current Biology 17, 922–931. 10.1016/j.cub.2007.05.018.

67. Engelsdorf, T., and Hamann, T. (2014). An update on receptor-like kinase involvement in the maintenance of plant cell wall integrity. Annals of Botany 114, 1339–1347. 10.1093/aob/mcu043.

68. Denness, L., McKenna, J.F., Segonzac, C., Wormit, A., Madhou, P., Bennett, M., Mansfield, J., Zipfel, C., and Hamann, T. (2011). Cell Wall Damage-Induced Lignin Biosynthesis Is Regulated by a Reactive Oxygen Species- and Jasmonic Acid-Dependent Process in Arabidopsis. Plant Physiology 156, 1364–1374. 10.1104/pp.111.175737.

69. Burda, I., Brauns, F., Shipman, A., Shapland, E., Hong, L., and Roeder, A. (2025). ROS inhibits microtubule dynamics and cell growth heterogeneity during Arabidopsis sepal morphogenesis. Preprint at Developmental Biology, 10.1101/2025.05.09.653143 https://doi.org/10.1101/2025.05.09.653143.

70. Gorgues, L., Smokvarska, M., Mercier, C., Igisch, C.P., Crabos, A., Dongois, A., Bayle, V., Fiche, J.-B., Nacry, P., Nollmann, M., et al. (2025). GEF14 acts as a specific activator of the plant osmotic signaling pathway by controlling ROP6 nanodomain formation. EMBO Rep 26, 2146–2165. 10.1038/s44319-025-00412-w.

71. Lü, S., Song, T., Kosma, D.K., Parsons, E.P., Rowland, O., and Jenks, M.A. (2009). Arabidopsis *CER8* encodes LONG□CHAIN ACYL□COA SYNTHETASE 1 (LACS1) that has overlapping functions with LACS2 in plant wax and cutin synthesis. The Plant Journal 59, 553–564. 10.1111/j.1365-313X.2009.03892.x.

72. Le Hir, R., Sorin, C., Chakraborti, D., Moritz, T., Schaller, H., Tellier, F., Robert, S., Morin, H., Bako, L., and Bellini, C. (2013). ABCG 9, ABCG 11 and ABCG 14 ABC transporters are required for vascular development in A rabidopsis. The Plant Journal 76, 811–824. 10.1111/tpj.12334.

73. Kleinboelting, N., Huep, G., Kloetgen, A., Viehoever, P., and Weisshaar, B. (2012). GABI-Kat SimpleSearch: new features of the Arabidopsis thaliana T-DNA mutant database. Nucleic Acids Research 40, D1211–D1215. 10.1093/nar/gkr1047.

74. Cutler, S.R., Ehrhardt, D.W., Griffitts, J.S., and Somerville, C.R. (2000). Random GFP∷cDNA fusions enable visualization of subcellular structures in cells of *Arabidopsis* at a high frequency. Proc. Natl. Acad. Sci. U.S.A. 97, 3718–3723. 10.1073/pnas.97.7.3718.

75. Marc, J., Granger, C.L., Brincat, J., Fisher, D.D., Kao, T., McCubbin, A.G., and Cyr, R.J. (1998). A *GFP–MAP4* Reporter Gene for Visualizing Cortical Microtubule Rearrangements in Living Epidermal Cells. Plant Cell 10, 1927–1939. 10.1105/tpc.10.11.1927.

76. Graumann, K., Runions, J., and Evans, D.E. (2010). Characterization of SUN-domain proteins at the higher plant nuclear envelope. The Plant Journal 61, 134–144. 10.1111/j.1365-313X.2009.04038.x.

77. Mouille, G., Ralet, M., Cavelier, C., Eland, C., Effroy, D., Hématy, K., McCartney, L., Truong, H.N., Gaudon, V., Thibault, J., et al. (2007). Homogalacturonan synthesis in *Arabidopsis thaliana* requires a Golgi□localized protein with a putative methyltransferase domain. The Plant Journal 50, 605–614. 10.1111/j.1365-313X.2007.03086.x.

78. Mølhøj, M., Verma, R., and Reiter, W.-D. (2004). The Biosynthesis of D -Galacturonate in Plants. Functional Cloning and Characterization of a Membrane-Anchored UDP- D - Glucuronate 4-Epimerase from Arabidopsis. Plant Physiology 135, 1221–1230. 10.1104/pp.104.043745.

79. Torres, M.A., Dangl, J.L., and Jones, J.D.G. (2002). *Arabidopsis* gp91^phox^ homologues *AtrbohD* and *AtrbohF* are required for accumulation of reactive oxygen intermediates in the plant defense response. Proc. Natl. Acad. Sci. U.S.A. 99, 517–522. 10.1073/pnas.012452499.

80. Alonso, J.M., Stepanova, A.N., Leisse, T.J., Kim, C.J., Chen, H., Shinn, P., Stevenson, D.K., Zimmerman, J., Barajas, P., Cheuk, R., et al. (2003). Genome-Wide Insertional Mutagenesis of *Arabidopsis thaliana*. Science 301, 653–657. 10.1126/science.1086391.

81. Edwards, K., Johnstone, C., and Thompson, C. (1991). A simple and rapid method for the preparation of plant genomic DNA for PCR analysis. Nucl Acids Res 19, 1349–1349. 10.1093/nar/19.6.1349.

82. Lampropoulos, A., Sutikovic, Z., Wenzl, C., Maegele, I., Lohmann, J.U., and Forner, J. (2013). GreenGate - A Novel, Versatile, and Efficient Cloning System for Plant Transgenesis. PLoS ONE 8, e83043. 10.1371/journal.pone.0083043.

83. Clough, S.J., and Bent, A.F. (1998). Floral dip: a simplified method for *Agrobacterium* Dmediated transformation of *Arabidopsis thaliana* . The Plant Journal 16, 735–743. 10.1046/j.1365-313x.1998.00343.x.

84. Daudi, A., and O’Brien, J.A. (2012). Detection of Hydrogen Peroxide by DAB Staining in Arabidopsis Leaves. Bio Protoc 2, e263.

85. Schindelin, J., Arganda-Carreras, I., Frise, E., Kaynig, V., Longair, M., Pietzsch, T., Preibisch, S., Rueden, C., Saalfeld, S., Schmid, B., et al. (2012). Fiji: an open-source platform for biological-image analysis. Nat Methods 9, 676–682. 10.1038/nmeth.2019.

86. Barberon, M., Vermeer, J.E.M., De Bellis, D., Wang, P., Naseer, S., Andersen, T.G., Humbel, B.M., Nawrath, C., Takano, J., Salt, D.E., et al. (2016). Adaptation of Root Function by Nutrient-Induced Plasticity of Endodermal Differentiation. Cell 164, 447–459. 10.1016/j.cell.2015.12.021.

